# Interplay between particle size and microbial ecology in the gut microbiome

**DOI:** 10.1101/2024.04.26.591376

**Authors:** Jeffrey Letourneau, Verónica M Carrion, Sharon Jiang, Olivia W Osborne, Zachary C Holmes, Aiden Fox, Piper Epstein, Chin Yee Tan, Michelle Kirtley, Neeraj K Surana, Lawrence A David

## Abstract

Physical particles can serve as critical abiotic factors that structure the ecology of microbial communities. For non-human vertebrate gut microbiomes, fecal particle size (FPS) has been known to be shaped by chewing efficiency and diet. However, little is known about what drives FPS in the human gut. Here, we analyzed FPS by laser diffraction across a total of 76 individuals and found FPS to be strongly individualized. Surprisingly, a behavioral intervention with 41 volunteers designed to increase chewing efficiency did not impact FPS. Dietary patterns could also not be associated with FPS. Instead, we found evidence that mammalian and human gut microbiomes shaped FPS. Fecal samples from germ-free and antibiotic-treated mice exhibited increased FPS relative to colonized mice. In humans, markers of longer transit time were correlated with smaller FPS. Gut microbiota diversity and composition were also associated with FPS. Finally, *ex vivo* culture experiments using human fecal microbiota from distinct donors showed that differences in microbiota community composition can drive variation in particle size. Together, our results support an ecological model in which the human gut microbiome plays a key role in reducing the size of food particles during digestion, and that the microbiomes of individuals vary in this capacity. These new insights also suggest FPS in humans to be governed by processes beyond those found in other mammals and emphasize the importance of gut microbiota in shaping their own abiotic environment.

## INTRODUCTION

The size of abiotic particles in the environment can influence microbial community composition via differences in nutrient availability and by the physical separation of microbes into sub-communities. In marine ecosystems, particle-associated communities undergo waves of succession, with primary degraders colonizing first, followed by secondary consumers of byproducts produced by the primary degraders^1,2^. In soil, which is often classified by particle size distributions^3^, smaller particle fractions are associated with the greatest diversity of both bacteria and fungi^4^. Individual microbes may have affinities for specific particle sizes^5^, and particle size fractions have been associated with differences in metabolic function^6,7^. The division of communities onto discrete particles may also be viewed as a form of spatial partitioning, an important factor in community assembly and diversity^8^. At the same time, the physical breakdown of particles is a means through which microbes shape their environments^9–11^.

For vertebrate gut microbiomes, fecal particle size (FPS) has been studied for its impact on community structure and composition in the distal colon, where evidence of particle-dependent community assembly and metabolism has been observed. Broadly, results have supported a model in which FPS is principally driven by chewing efficiency^12–16^. Reptiles, which do not chew their food, have higher FPS than mammals of equivalent body mass^13^, and proboscis monkeys, which ruminate, have smaller FPS than similar-sized monkeys, considering the known relationship between FPS and weight across species^12^. Diet particle size has also been shown to influence the gut microbiome, particularly in the feeding of livestock^17^. In sheep, reduced feed particle size has been shown to cause increases in the abundance of certain fiber-degrading taxa, and in overall diversity, providing *in vivo* support for the hypothesis that intake particle size influences microbiome composition and metabolism^18^. In cattle, altering feed particle size has been shown to influence microbiome composition and short-chain fatty acid (SCFA) concentrations^19^. Gut microbes have been thought to play a direct role in particle size reduction only in cases where gastrointestinal transit time is exceptionally long, such as in the dugong^12,20^.

In contrast to our understanding for how mastication efficiency and diet shape FPS in most mammals, little is known about the factors shaping FPS in the human gut. Particles encountered by human gut microbes have been shown to be food-derived, including structures that escape digestion by human enzymes, such as plant cell walls and resistant starch granules, as well as host compounds like mucin, and microbes do adhere to such particles^21,22^. Chewing has been shown to reduce *intake* food particle size in humans^23–27^, but these prior studies only evaluated particle size by analyzing chewed food that was then expectorated instead of measuring it in feces. Rather than focus on which factors govern FPS, most prior work has instead focused on how FPS governs gut microbial ecology. Particle-associated gut microbes have been shown to differ taxonomically from those found in the liquid phase^28^. Smaller particle sizes are also known to promote enhanced production of SCFA, likely through increased surface area of accessible nutrients^29–33^, though some reports have found coarser particles may actually deliver more butyrate to the distal colon^30^. Ultimately, the factors shaping FPS in humans remain unstudied, and the assumption that gut microbes play a negligible role in particle breakdown in humans remains untested.

Given the influence of particle size on the assembly and metabolism of microbial communities, we sought to understand the factors shaping FPS in the human gut. We hypothesized that mastication and diet would shape the distribution of FPS across individuals. Based on prior work among vertebrates, we expected that lower FPS could be induced by increasing chewing efficiency or through select dietary patterns. In essence, we sought to understand whether modifiable behavioral factors could influence the physical particulate environment in the gut, and by extension, microbiome ecology and metabolism (Fig. 1).

**Figure 1.**
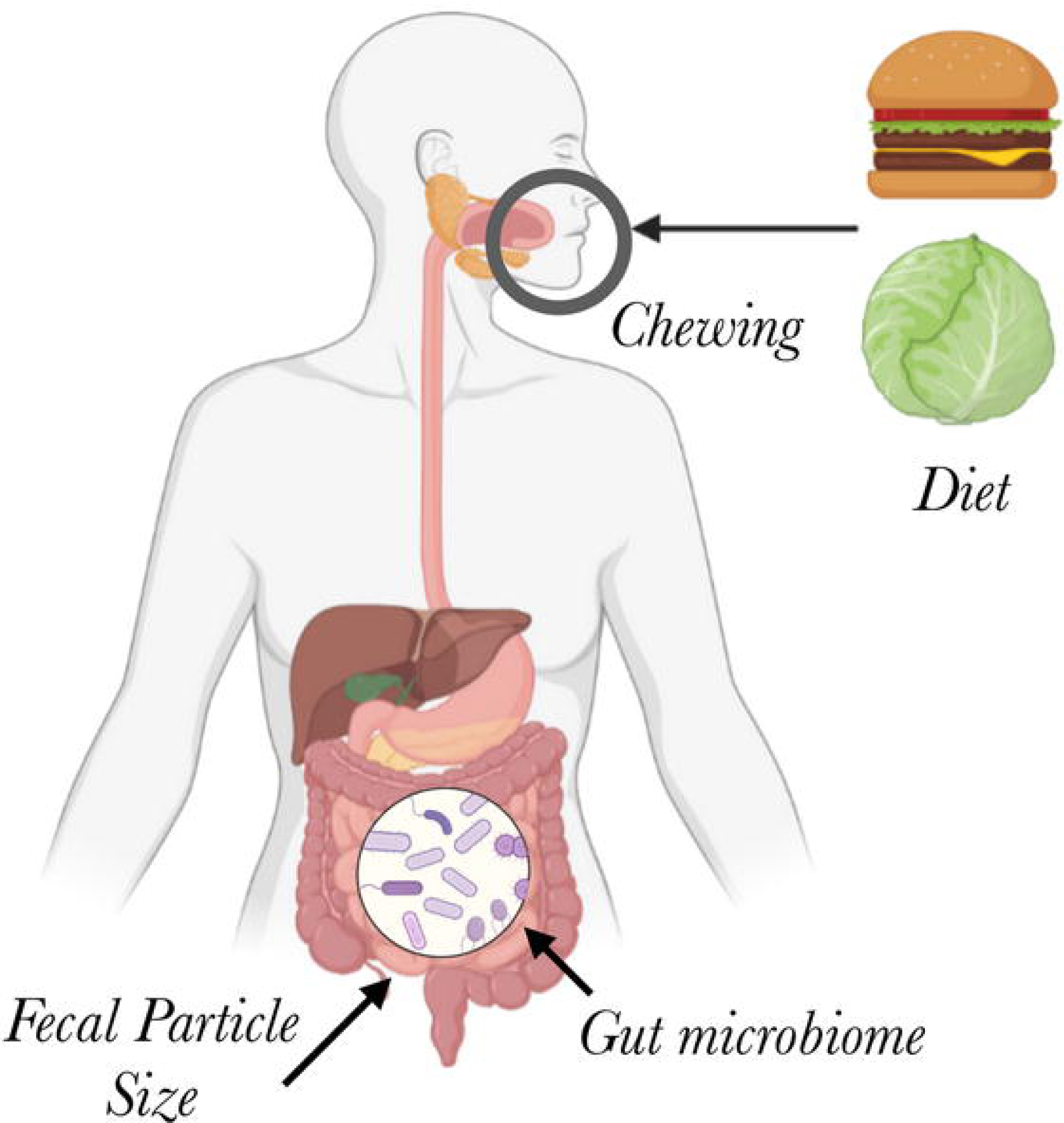
Factors potentially influencing fecal particle size. We considered in this study a model in which FPS is driven by either dietary patterns, the degree to which foods were chewed, or gut microbial metabolism.

## RESULTS

### Fecal particle size is individualized

To assess the influence of diet and chewing, we enrolled 41 healthy adults in a fixed-order within-subjects study to test the effects of chewing and habitual macronutrient intake on FPS over a two-week study period (Cohort 1; Fig. S1). Participants consumed their typical diets on week 1 (baseline week) and were asked to chew every bite of food more thoroughly on week 2 (chewing week), by chewing each bite 30-40 times or until the food had reached roughly an applesauce consistency. We also re-analyzed samples and behavioral data from a cohort of 35 healthy adults who we previously enrolled in a study of prebiotics and cognitive performance^34^ (Cohort 2). We surveyed macronutrient intake using the Diet History Questionnaire III (DHQ3) across both cohorts, an approach that has previously revealed associations between dietary patterns and the gut microbiome in prior studies carried out by our group^34,35^, and we analyzed stool samples for fecal particle size and microbiome composition.

We used laser diffraction to analyze fecal particle size across the two cohorts. The average median value across all participants was 44.5 µm, and the average volume-weighted particle size distribution showed a peak at 0.872 µm and a second, higher peak at 111 µm, with only one sample of 186 analyzed (0.54%) having any particles detectable above 1000 µm (Fig. 2B).

**Figure 2.**
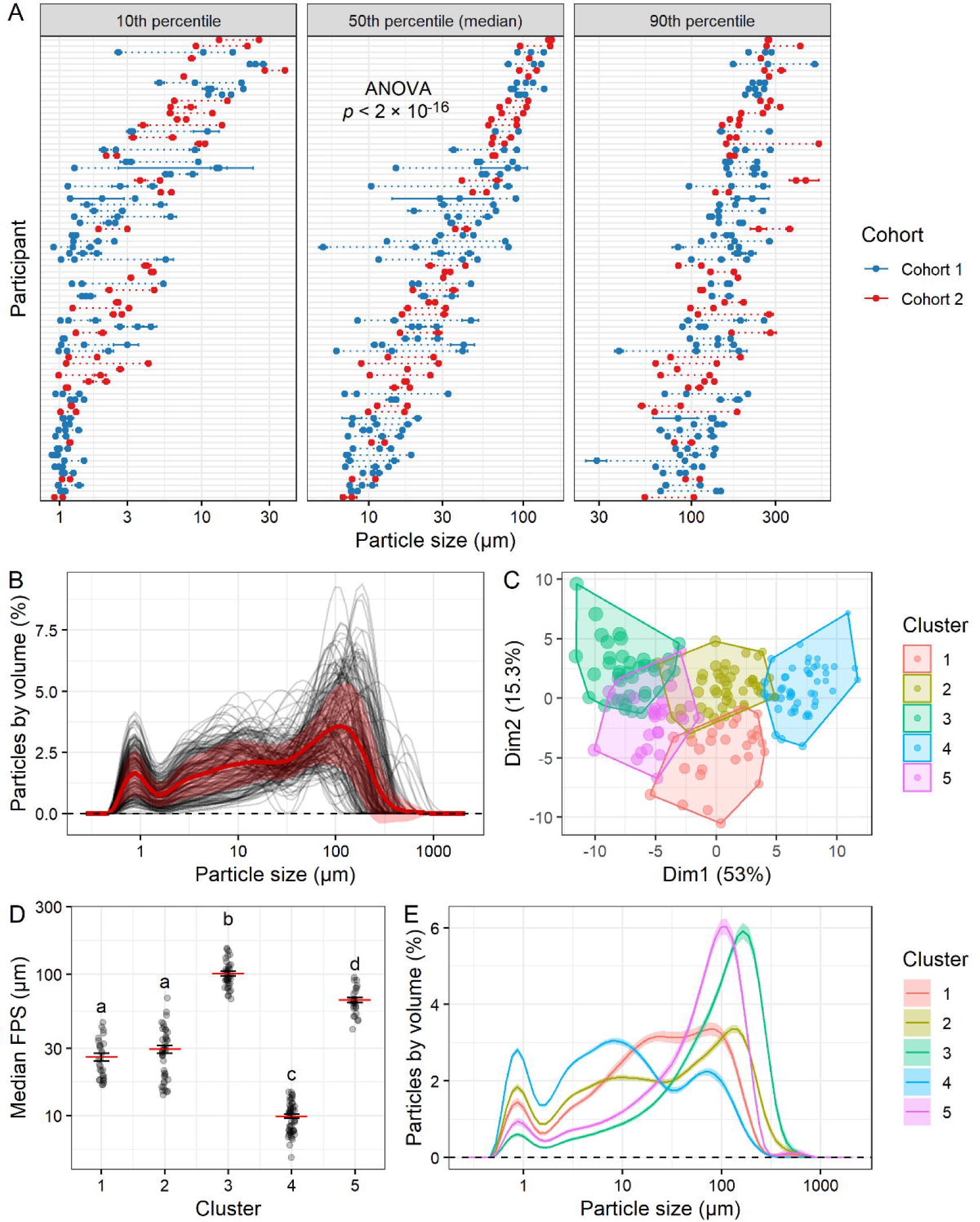
Fecal particle size varies across individuals. **A**, Summary metrics of FPS distributions in all baseline samples by participant, arranged by individual average median value, with result of ANOVA on participant shown. Dotted lines connect samples from the same participant, and error bars denote standard error of multiple laser diffraction measurements of the same sample. **B**, Volume-weighted particle size distributions for the same sample set, with mean and standard deviation plotted in red. **C**, Principal component visualization using *k*-means clustering. Point size corresponds to median FPS of each sample. **D**, Median FPS by cluster, with mean and standard error shown. ANOVA (cluster + participant) cluster *p* < 2 × 10^−16^, participant *p* = 0.00034; differences by Tukey HSD test shown. **E**, Mean and standard error particle size distribution curves based on the five clustered sets. **A-E**, (*n* = 186 samples from 76 participants; 41 in Cohort 1 and 35 in Cohort 2.)

We noticed our median FPS was lower than the FPS reported among primates of comparable mass to humans (*Pan troglodytes* at 52 kg, *Pongo pygmaeus* at 60 kg, and *Gorilla gorilla* at 98 kg) at about 3 mm (2.9, 2.4, and 3.6 mm, respectively)^55^. We therefore analyzed fecal samples from three species of lemurs, two of which previously had FPS measured in the literature^55^. We found that *Varecia variegata* (*n* = 6) had a median FPS averaging 101 µm (in contrast to 2,114 µm previously described^55^) and measured the median FPS of *Lemur catta* (*n* = 6) at an average of 65 µm (compared to 1914 µm previously described^55^; Fig. S2). Furthermore, we analyzed mouse fecal pellets (*n* = 10 specific pathogen-free [SPF] mice) and found they had a median FPS averaging 111 µm (Fig. S2) compared to a previously reported value of 209 µm (for a single mouse)^55^. These results suggest that, compared to prior work using wet sieving methods (e.g. smallest sieve size commonly of 63 or 125 µm^12–14,55^), our technique may have increased resolution of smaller fecal particles.

Independent of the range of FPS values, we observed a substantial amount of inter-individual variation in FPS. Analyzing the baseline (i.e. pre-intervention) samples from these two cohorts, we found that participant was a significant explanatory variable both in terms of median FPS (ANOVA by participant *p* < 2 × 10^−16^; Fig. 2A) and the full particle size distribution (PERMANOVA by participant *p* < 0.0001; Fig. S3A). These results support the hypothesis that FPS is an individualized metric which may be dependent on host-specific factors.

To more easily visualize the common forms of particle size distributions, we performed *k*-means clustering, partitioning the data into five clusters (Figs. 2C-D, S3B-D). These clusters were partially distinguished by median FPS (Fig. 2D), but also by the shape of the distribution, particularly in comparing clusters 1 and 2 (Fig. 2E). Cluster 3 had the highest median FPS, followed by Cluster 5; Clusters 1 and 2 had intermediate median FPS, with Cluster 1 defined by a tighter distribution, whereas Cluster 2 was skewed more towards both the high and low ends; Cluster 4 had the lowest median FPS (Fig. 2D-E). In the majority of cases, samples from the same individual were in the same cluster, or adjacent clusters (Fig. S3D).

### Origins of variation in particle size

To investigate the mechanism behind the particle size distributions we observed, we first considered the hypothesis that host processing of food by mastication might drive FPS, an association that the study design of Cohort 1 should support if the hypothesis is true (Fig. 1) Compliance in the chewing intervention in Cohort 1 as assessed by self-reported survey responses was as desired; participants remembered to chew as requested for 83.3 ± 13.6% (mean ± sd) of solid meals consumed during the chewing week. Participants also reported significantly higher chewing thoroughness (from a mean of 5.3 to 8.5 on a 1-10 scale), average number of chews per bite (from 8.5 to 27.8), and meal duration (from 18.3 to 28.5 min) during the chewing week compared to their baseline (linear mixed model *p* < 0.001; Fig. 3A). Additional evidence for the effectiveness of our intervention was that a majority of participants (22/39, 56.4%) reported that the chewing intervention reduced the amount of food they consumed when asked as part of the post-intervention survey (Fig. S4C), consistent with past research indicating that additional chewing leads to increased satiety and reduced food intake^38^. Yet, surprisingly, increased chewing in Cohort 1 did not lead to a reduction in median FPS as expected (linear mixed model by week*day_of_week NS for all terms, *p* = 0.22 for week; Figs. 3B, S5). There was also no difference in terms of overall particle size distributions (PERMANOVA *p* = 0.70; Fig. S6A). Our findings therefore do not support the hypothesis that chewing efficiency regulates FPS in humans.

**Figure 3.**
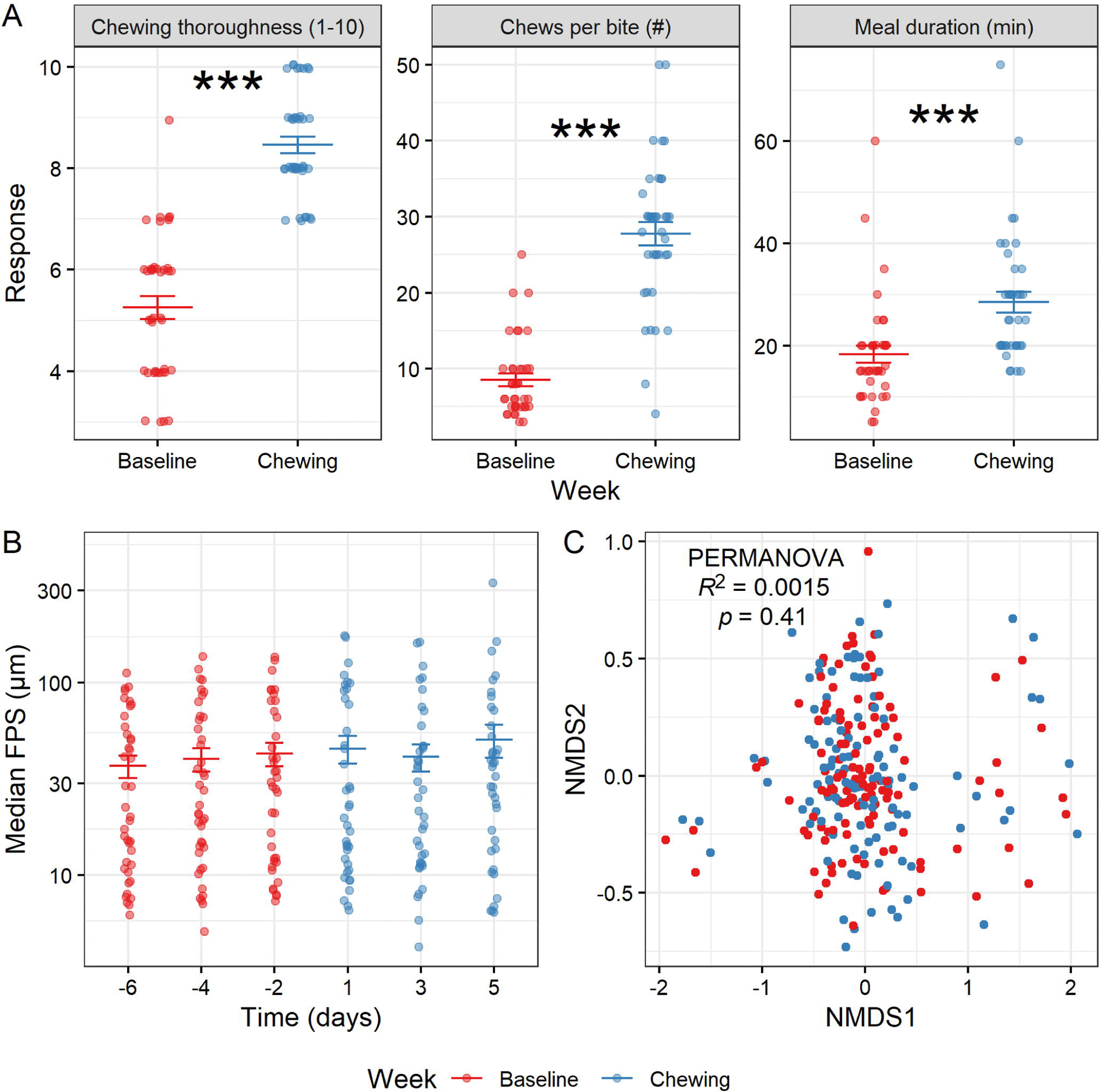
Results of chewing intervention study. **A**, Self-reported compliance assessed by comparison of pre-study survey responses relating to typical chewing efficiency, typical number of chews per bite, and typical meal duration to these same metrics assessed for the intervention week. Linear mixed model (week as fixed effect, participant as random effect) shown. (*n* = 39 participants). **B**, Median FPS by time point in chewing study. Linear mixed model (time as categorical fixed effect, participant as random effect) with day −6 as intercept shown. LMM by week*day_of_week *p* > 0.05 for all terms. (*n* = 41 participants). **A-B**, Mean and standard error shown. * *p* < 0.05, ** *p* < 0.01, *** *p* < 0.001. **C**, NMDS plot of community composition by 16S rRNA gene sequencing, with results of PERMANOVA on week (participant as strata) shown. PERMANOVA on day *R*^2^ = 0.0054, *p* = 0.95. (*n* = 41 participants).

While the chewing intervention in Cohort 1 was not associated with FPS, we note that variation in chewing could be linked to changes in individuals’ microbiome structure and function. We analyzed individualized alterations to community composition by calculating beta-dissimilarity relative to each individual’s own baseline and found significant compositional changes associated with the treatment week (Fig. S6C-E). Alpha diversity and overall inter-individual variation in community composition, though, did not change across the intervention (Fig. S6B; Fig. 3C). These findings together suggest the possibility of individualized community shifts, but not a case of the same taxa increasing across most participants. We also observed a difference in fecal SCFA content; total SCFA was significantly decreased by day 1 and in the chewing week overall (LMM *p* < 0.05; Fig. S6F), driven most apparently by a decrease in propionate (Fig. S6G), with proportions of isobutyrate and isovalerate significantly increased (Fig. S6H). The observed decrease in SCFA was not related to alterations in diet or host metabolism that occurred as a result of the chewing intervention, as we detected no differences in macronutrient proportions as a fraction of total calories between the two study weeks by ASA24 dietary recall survey (Fig. S4A-B). Thus, some changes to gut microbial ecology occurred during the chewing intervention, but these changes appeared independent of shifts in FPS.

We next investigated the hypothesis that diet is a driver of FPS in humans. We saw no evidence for such an association in either of our cohorts. There was no detectable relationship between macronutrients and FPS in Cohorts 1 and 2 (Fig. S7A-B). When we performed FoodSeq, a genomic technique used to identify residual food DNA in stool^37^, on size-fractionated samples from Cohort 2, we detected dietary plant DNA across all particle size fractions (Fig. S8D-E). However, we found neither an association between diet and particle size nor one between dietary plant species composition in stool samples and FPS (PERMANOVA *p* > 0.05; Fig. S7C). Therefore, our data did not support the hypothesis that intake of specific macronutrients or plant species are major drivers of FPS in humans.

### Gut microbes play a role in particle breakdown

Since neither mastication nor diet could be linked to inter-individual variation in FPS, we next considered the hypothesis that FPS is mediated by microbiome activity. Prior studies have found chewing and diet to be major drivers of FPS rather than the microbiome^12–16^. However, *in vitro* evidence has shown degradation of food particles by microbes^33^, and *in vivo* studies have shown that microbes may play a role in FPS over sufficiently long timescales of digestion^20^ and even on shorter time scales in the context of fine particles^39^.

We first confirmed that indeed, gut microbes could degrade particles in the mammalian gut. We analyzed two *in vivo* sample sets in mice, reasoning that if microbes played a role in particle size breakdown, then germ-free (GF) mice would have a higher FPS than colonized counterparts, and that reduction in gut microbial load within C57BL/6 mice with antibiotics would lead to an increase in FPS. As expected, we found that GF mice (*n* = 10) had greater median FPS than SPF mice (*n* = 12) (linear model *p* = 0.0082; Fig. 4A-B). Mice treated with the antibiotic imipenem (*n* = 5 cages of 3 mice each) also exhibited a significant increase in FPS as anticipated (linear mixed model with treatment day as categorical fixed effect and cage as random effect *p* = 0.042 on day 1 and *p* = 0.00046 on day 2; Fig. 4C-D). Both GF and antibiotic-treated mice exhibited a clear decrease in the 0.872 µm micron peak (Fig. 4B,D). This apparent decrease, combined with the specific size, suggests that this peak corresponds to free bacteria. Therefore, to ensure that we were not simply capturing differences due to differences in total bacterial biomass, we also analyzed our data omitting particles < 2 µm and still found that GF and antibiotic-treated mice had increased FPS (Fig. S9B-E).

**Figure 4.**
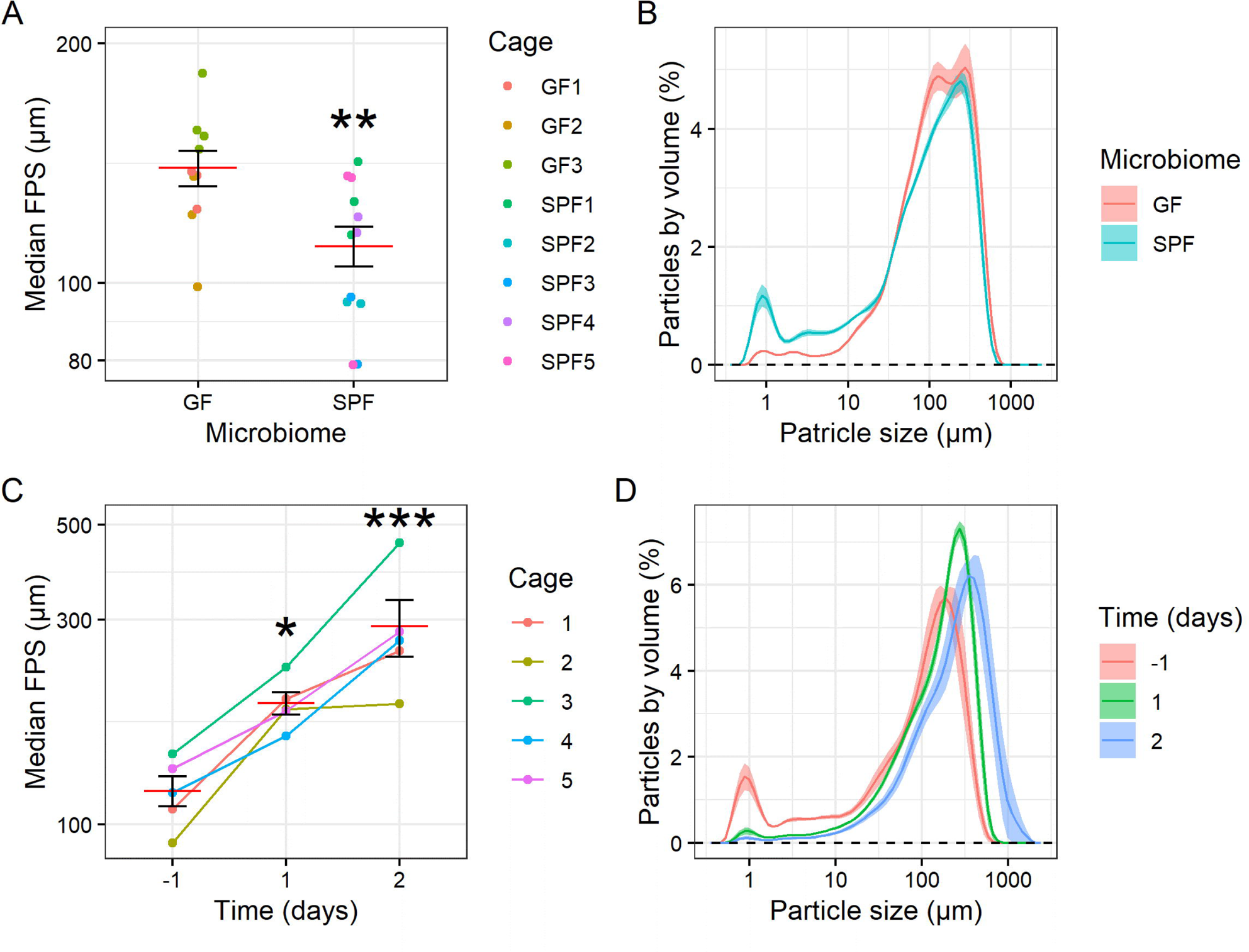
State of gut microbiome influences FPS in mice. **A**, Median FPS in germ-free (GF) and specific-pathogen-free (SPF) mice collected from individual mice. (Linear model with GF as intercept; *n*LJ=LJ10 GF mice and 12 SPF mice.) **B**, Particle size distributions for (**A**). **C**, Median FPS for mouse fecal samples collected by cage, with imipenem antibiotic treatment begun on day 0. Linear mixed model (day as categorical fixed effect, cage as random effect) with D-1 as intercept shown. (*n*LJ=LJ5 cages.) **D**, Particle size distributions for (**C**). Mean and standard error plotted. * *p* < 0.05, ** *p* < 0.01, *** *p* < 0.001.

Since gut microbial load impacted FPS in mice, we next reasoned that FPS could be shaped by the length of time microbes had to interact with particles. Studies have indicated that stool consistency as measured by Bristol score is correlated with intestinal transit time^40^, so we analyzed the relationship between self-reported typical Bristol score and baseline FPS across both cohorts (Fig. S10C) and between chewing week Bristol score with FPS in Cohort 1 (Fig. S10D). While there was a non-significant trend towards higher Bristol score being associated with higher median FPS, the low numbers of individuals reporting Bristol scores outside the 3-4 range did not enable a definitive conclusion (Fig. S10C-D). Therefore, for a higher resolution alternative, we measured the moisture content of Cohort 1 stool samples by lyophilizing samples and comparing dry mass to the original sample mass. Fecal moisture content is known to decrease as water is absorbed by intestinal cells, and may therefore serve as an alternative and more granular measure of gastrointestinal transit time. Consistent with a model in which longer microbial exposure would decrease FPS, we found a significant difference in moisture content by particle size distribution cluster (ANOVA *p* = 1.6 × 10^−6^). The clusters with higher FPS also had higher moisture content, and thus an inferred lower transit time, whereas lower FPS clusters had lower moisture content (Fig. 5A). These results suggest that gastrointestinal transit time plays a role in FPS in the mammalian gut.

**Figure 5.**
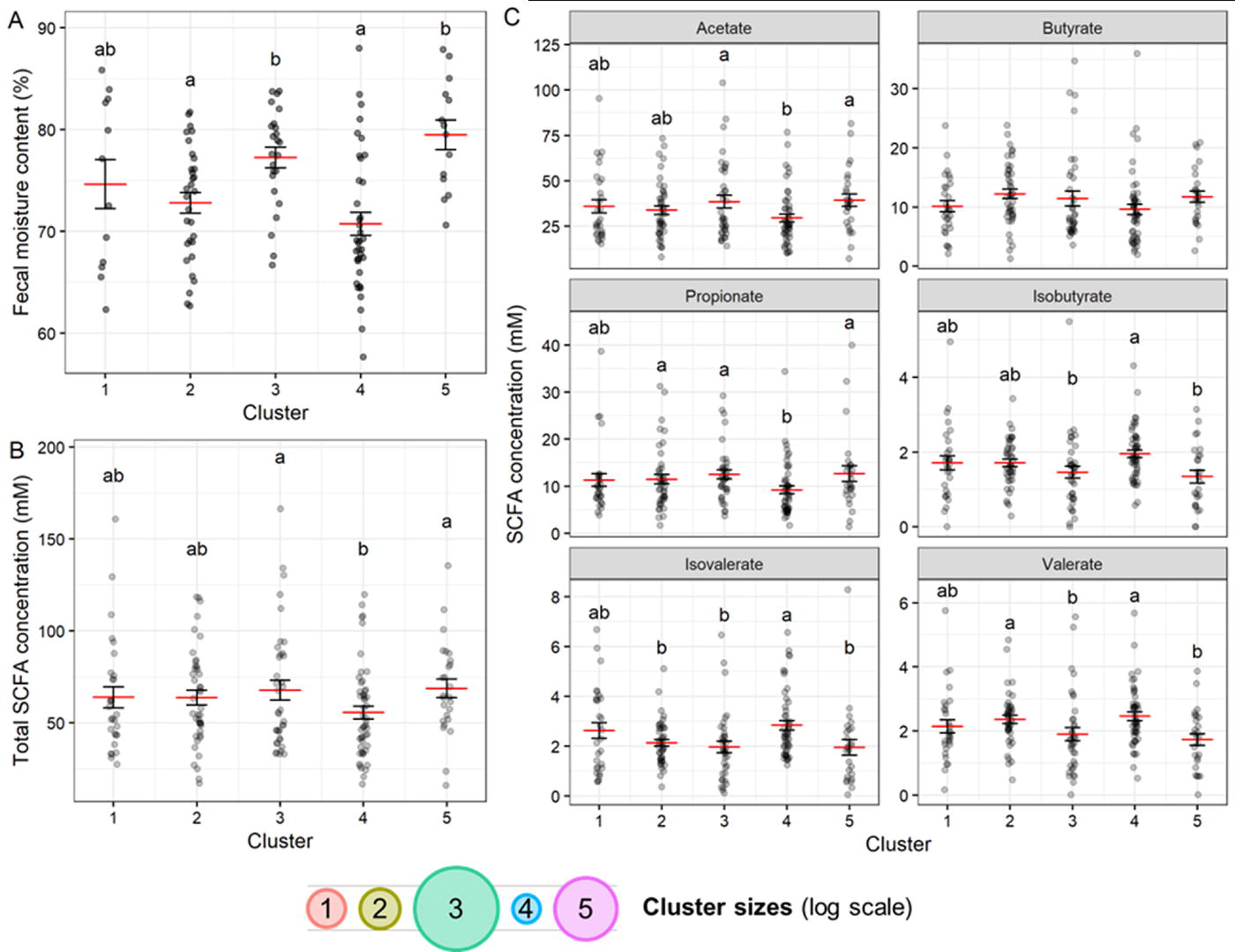
Fecal particle size relates to short-chain fatty acid and moisture content. **A**, Moisture content of Cohort 1 stool samples by particle size distribution cluster. ANOVA (cluster + participant) cluster p = 1.6 × 10-6, participant p = 0.0035; letters indicate significantly different groups by Tukey HSD test. (*n* = 118 samples). **B**, Total SCFA concentrations for all baseline samples by particle size distribution cluster. **C**, Concentrations of individual SCFAs by cluster. **A-B**, ANOVAs (cluster + participant) significant for both variables for total and all individual SCFAs except butyrate. (*n* = 185 samples). **A-C**, Mean and standard error shown. Where effect of cluster p < 0.05 by ANOVA, letters indicate significantly different groups by Tukey HSD test.

We hypothesized that the decrease in FPS observed with longer transit times could be the result of increased breakdown of fecal particles. Longer microbial degradation of food particles could also lead to shifts in substrate consumption as primary substrates are consumed; for example, we would predict increases in branched chain fatty acids (as the community shifts from saccharolytic to proteolytic fermentation)^41^. Finally, as transit time increases, we could anticipate two potential impacts on total SCFA levels. With longer transit SCFAs could increasingly be taken up by host intestinal cells^42^, meaning smaller FPS could also be associated with a reduction in total fecal SCFA content. Yet, others have reported that smaller particle sizes should increase SCFA metabolism^30,31^, potentially through increasing the surface area of available nutrients^30^. Consistent with these reports, *in vitro* experiments in which we modulated the size of wheat bran particles demonstrated that smaller wheat bran particles elicited greater SCFA production (Fig. S11A).

To assess these predictions in our human samples, we profiled the SCFA content of stool samples in both cohorts. We found that total SCFA content varied with particle size cluster (ANOVA *p* = 0.020 for cluster and 3.6 × 10^−10^ for participant). The clusters with larger median particle sizes (Clusters 3 and 5), which also were associated with higher moisture content, had higher fecal SCFA concentrations; the cluster with the smallest particle sizes (Cluster 4), which was associated with lower moisture content, had the lowest median SCFA concentration (Fig. 5B-C). These findings suggest that the *in vivo* effects of longer transit time on increased SCFA absorption outweigh the ecological effects of having more surface area for bacterial metabolism.

Overall variation in SCFA levels were primarily driven by differences in the concentrations of acetate and propionate in stool samples, whereas the concentrations of the branched chain amino acids isobutyrate, isovalerate, and valerate tended to exhibit the opposite trend, with concentrations higher in the cluster with the smallest FPS (Cluster 4; Fig. 5C).

### Community composition relates to FPS

If gut bacterial metabolism influences FPS, we would expect that different microbiota compositions, which have inherently different metabolic capacities, would be associated with specific particle size distributions that are the result of these metabolic processes. To test this hypothesis, we analyzed FPS data in the context of microbiota composition features using 16S rRNA gene sequencing. We found that across individuals, median FPS was negatively correlated with alpha diversity both in observed ASVs (Spearman’s rho = −0.64, *p* = 5.6 × 10^−10^) and Shannon diversity (Spearman’s rho = −0.59, *p* = 3.0 × 10^−8^; Fig. 6A). This trend of smaller FPS samples being associated with greater diversity is consistent with previous findings in soil microbiomes^4^. Differences in read depth did not explain this relationship (Spearman’s correlation *p* > 0.05; Fig. S12A-B). Moreover, particle size distribution clusters significantly differed by alpha diversity (ANOVA *p* < 0.05; Fig. 6B).

**Figure 6.**
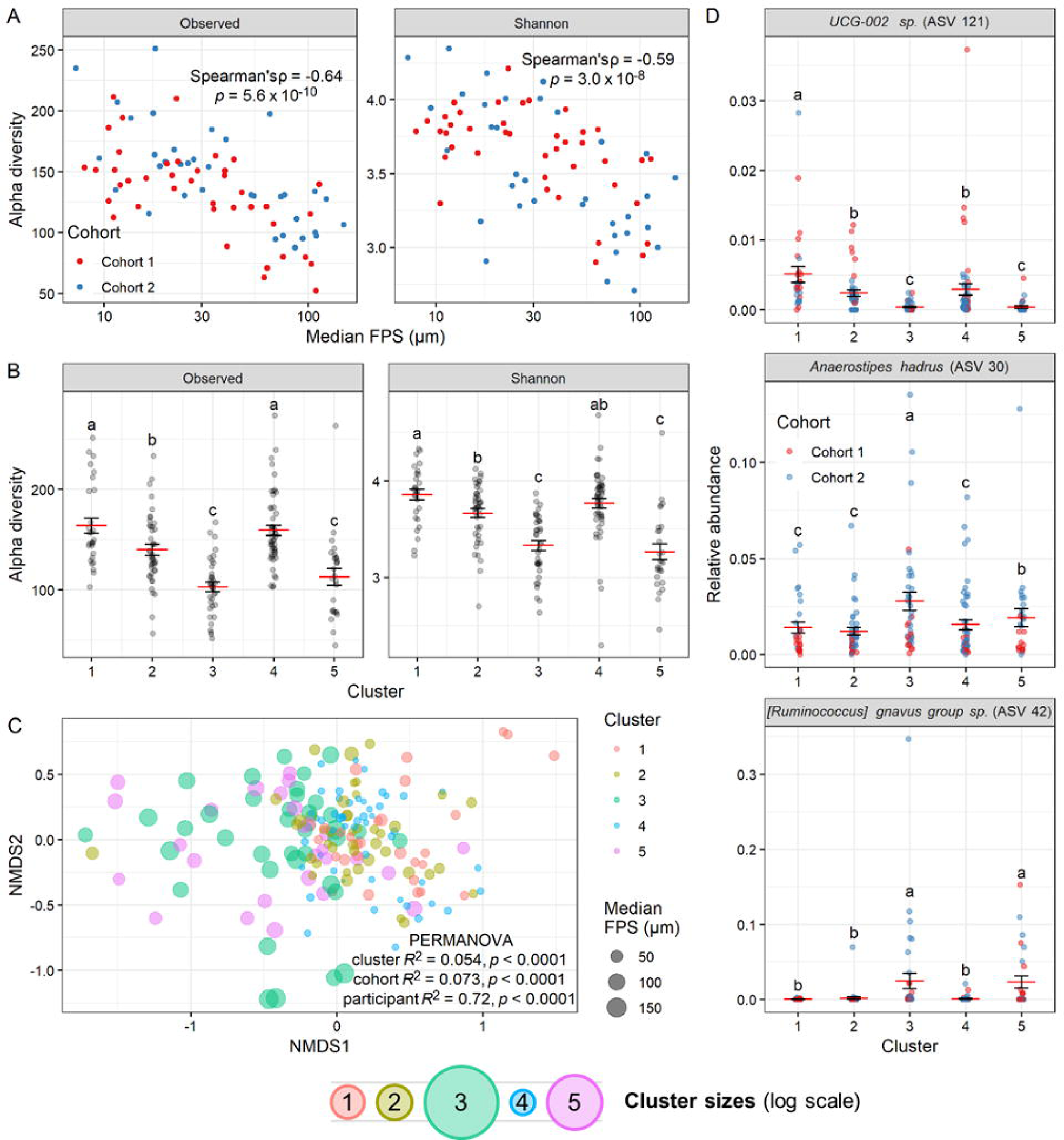
Fecal particle size relates to gut microbiome composition profiled by 16S rRNA gene amplicon sequencing. **A**, Correlation between median FPS averaged within each participant and alpha diversity by observed ASVs and Shannon index. Spearman correlation test statistics shown. (*n* = 76 participants; 41 in Cohort 1 and 35 in Cohort 2.) **B**, Alpha diversity metrics for all individual samples by particle size distribution cluster. ANOVAs (cluster + participant) significant for both variables for both diversity metrics; letters indicate significantly different (*p* < 0.05) groups by Tukey HSD test. **C**, NMDS ordination plot of 16S data for all samples, with results of PERMANOVA (cluster + cohort + participant) shown. **D**, Relative abundances by cluster for top three ASVs by lowest ALDEx2 Kruskal-Wallis test *p*-value assessing compositional differences by cluster. Letters indicate significantly different groups by Tukey HSD test on CLR-transformed count data. **B,D**, Mean and standard error plotted. **B-D**, (*n* = 185 samples).

Examining taxonomic composition, we found significant differences both by median FPS (Fig. S13A) and cluster (Fig. 6C). While we also found significant differences by cohort (Fig. S13A), which may be due to the use of different DNA extraction kits, we were still able to identify specific taxa differing by cluster by ALDEx2, with 29 of 298 analyzed ASVs (9.7%) having a Kruskal-Wallis *p*-value < 0.05 after multiple-comparisons correction (Fig. S13B). These included the Oscillospiraceae *UCG-002 sp.* (ASV 121), which was most abundant in Cluster 1, *Anaerostipes hadrus* (ASV 30), which was most abundant in Cluster 3, and *[Ruminococcus] gnavus group sp.* (ASV 42), which was most abundant in Clusters 3 and 5 (Fig. 6D). Taxa more abundant in high FPS samples were mostly members of Lachnospirales, and those more abundant in low FPS samples were mostly members of Oscillospirales (Fig. S13C). Members of the Oscillospiraceae family have previously been associated with constipation and disease states^43–45^.

Additionally, we observed evidence that distinct taxa were associated with specific particle sizes across individuals (Fig. S8A). We fractionated samples among 10 donors from Cohort 1 using sequential particle size filtration using a series of nylon mesh and paper filters, separating particles into four fractions of <2 µm, 2-11 µm, 11-100 µm, and >100 µm. We subsequently performed 16S rRNA sequencing on each size fraction. We observed higher relative abundance of two *Blautia* ASVs (Fig. S8B-C) in fractions containing smaller particles; members of this genus have previously been associated with colonization of small particles of wheat bran in the context of the human fecal microbiota^32^.

To directly test whether human gut microbiota composition impacts FPS, we performed an *in vitro* experiment with cultured microbial communities from 10 different human donors from Cohort 1. Each of these microbial communities was cultured with disinfected substrate stool from three distinct Cohort 1 donors (plus no-inoculation control) to determine whether differences in microbial composition among the 10 inoculation donors (from Cohort 2) affected the final particle size distribution of the substrate cultures. We found that donor identity had a consistent effect across substrates in terms of the final particle size distribution (PERMANOVA on inoculation donor with substrate donor as strata *p* = 0.0012, R^2^ = 0.18; Fig. S9A). In concert, these culture experiments and our analyses associating 16S rRNA composition with FPS size and clusters support the hypothesis that FPS is shaped by the specific composition of microbial taxa in the human gut.

## DISCUSSION

Our results support the overall hypotheses that human FPS is an individualized trait shaped by the gut microbiome as food matter transits through the intestinal tract, and that human FPS is not strongly associated with diet or mastication efficiency. Our data supports a model in which the more time gut microbes have in contact with particles, the smaller the size of those particles. Transit time may also explain why chewing resulted in decreased SCFA (Fig. S6F); increased chewing results in increased levels of glucagon-like peptide-1 (GLP-1)^24^, a hormone that inhibits gastric emptying^46^. Interestingly, a recent study found that individuals with shorter transit times unexpectedly had higher rates of energy extraction, further suggesting a key role of microbiome composition in energy extraction^47^.

One potential explanation for the lack of an association between human FPS and chewing or food choice, given the associations that have been shown in animals, are differences between the diets of wild animals and humans. Most human diets are composed of food that has been cooked, and studies suggest that thermal processing increases the energy gain from both starch- and meat-based foods^48^. The same physical changes that result in increased energy gain may also change the accessibility of dietary particles to microbial degradation. Modern food processing may also reduce particle size and food toughness^13,49–52^; millennia of plant domestication have produced crops that are easier to chew and contain lower fiber content than many wild varieties^53^, which is itself a form of processing^54^. Thus, the average diet consumed in our cohorts may contain particles that are too small to be sensitive to the effects of chewing and natural dietary variation. Furthermore, it is possible that the high heterogeneity of human diets, relative to those of other animals, might also add sufficient variance to mask effects of increased chewing.

We note that our findings that the microbiome is a key factor in shaping FPS neither rules out a potential impact of GI physiology nor feedbacks between FPS and gut microbial ecology. We were unable to measure, and thus exclude, a role for individual differences in mechanical (e.g. intestinal length or the force of peristalsis) or enzymatic processes in the stomach and small intestine^56^. Additionally, it is possible that FPS impacts on gut microbial ecology, such as the potential for SCFA production^29–33^, will in turn will shape gut microbiome structure. Such processes would be consistent with the associations we observe between microbial community composition and individual differences in FPS (Fig. S8A-C) and between altered substrate particle size and SCFA production (Fig. S11). Nevertheless, our culture-based experiments supporting a causal link between community composition and FPS are consistent with prior insights that the gut microbiome can shape their own abiotic environment^57^. FPS, like oxygen concentrations or redox conditions, may be an ecological factor that is both influenced by and shapes its ecology. Understanding the nuances of how microbes engage in particle adhesion and degradation may thus help us one day develop improved strategies for treating digestive disorders associated with gut microbial metabolism^58,59^.

Independent of the origins of FPS differences between individuals, our data indicating that FPS strongly correlates with gut microbial alpha diversity (Figs. 6A-B, S12C-D) and community composition (Figs. 6C-D, S13) argues that the well-known phenomenon of inter-individual microbiome variation could be related to inter-individual variation in FPS as well^60,61^. Notably, the inter-individual variation in FPS (PERMANOVA R^2^ = 0.74, Fig. S3A) is comparable in magnitude to inter-individual variation observed in 16S sequencing of human gut microbial communities (PERMANOVA R^2^ = 0.72, Fig. 6C) Overall, the results of our microbiome analysis suggest that host variability in microbiota composition and transit time could be related to a “signature” fecal particle size that is highly individualized. Considering the observed particle size distributions in the context of soil classification, the data spanned five categories, suggesting that the scale of particle size variability might be sufficient to correspond to differences in microbial ecology (Fig. S3E).

Such an association suggests FPS could serve as a useful biomarker. FPS has been used as a measure of impaired digestion in rainbow trout^62^, and could have similar applications as a metric of transit time or microbiota dysbiosis in humans. For example, slower transit time has been associated with disease conditions such as small intestinal bacterial overgrowth (SIBO) and increased risk of colon cancer^41^. Certain microbial community compositions have also been linked to increased energy harvest capacity in a manner that promotes obesity^63^. Yet, methods for tracking transit time (e.g. ingesting dyes and self-reporting its passage) may be considered inconvenient or unreliable and methods for tracking community composition (e.g. 16S rRNA sequencing) require familiarity with advanced genomic techniques. FPS analysis, by contrast, is objective and does not require access to advanced molecular techniques and instead only relies on the ability to homogenize fecal samples and access to particle size measurement by laser diffraction or other methods (e.g. sieving). Thus, FPS could provide a new, accessible, and informative biomarker of the compositional and metabolic state of the human gut microbiome.

## MATERIALS AND METHODS

### Participant recruitment and sample collection

Data from Cohort 2 constitutes a *post hoc* analysis of samples left over from a previously described^34^ cohort recruited primarily to test a link between the gut microbiome and aspects of cognition and behavior. The original study protocol was approved by the Duke Health Institutional Review Board (IRB) at Duke University under protocol number Pro00093322, and registered on ClinicalTrials.gov with the identified number NCT04055246.

Cohort 1 was recruited specifically to study the role of particle size in the gut microbiome; for this study, we recruited 41 healthy adult participants under Duke Health IRB protocol number Pro00108805 and ClinicalTrials.gov registration number NCT05091801 (Fig. S1). Participants were recruited by use of flyers on Duke University campus as well as electronic postings on DukeList (a university-internal classifieds website), a lab website, and ClinicalTrials.gov. All participants provided written informed consent via an electronic consent form (eConsent) prior to participation in the study protocol. Additional details for this cohort are discussed below.

All participants were at least 18 years old, and individuals were excluded from participation if they: had a colonoscopy or oral antibiotics within the past month; had a history or current diagnosis of irritable bowel syndrome, inflammatory bowel disease, type 2 diabetes, chronic kidney disease with decreased kidney function, intestinal obstruction, or untreated colorectal cancer; typically consumed more than one full meal per week in liquid form (smoothie, meal replacement drink, etc.); had any dental issues or other physical limitations that would prevent them from thoroughly chewing their food; or were already practicing mindfulness or dietary techniques involving increased chewing. Participants were 80.5% women (33/41), 17% men (7/41), and 2.4% (1/41) nonbinary. Participants were 51.2% white (21/41), 46.3% Asian (19/41), 4.9% Native American (2/41), 0% Black, 0% Native Hawaiian or other Pacific Islander, and 4.9% were another race not listed (2/41); additionally, 12.2% were Hispanic or Latino (5/41). Average age at time of enrollment was 26.4 ± 8.6 years (mean ± stdev). Most participants (80.5%; 33/41) were omnivores, 4.9% ate everything except red meat (2/41), 4.9% were vegetarian (2/41), 4.9% were vegetarian with the exception of fish (2/41), and 4.9% were vegan. Average weight was 63.5 ± 9.0 kg and average height was 166.5 ± 2.1 cm.

Participants provided stool samples three times per week on Monday, Wednesday, and Friday over a two-week study period. Samples were self-collected using polypropylene scoop-cap tubes (Globe Scientific), and participants were instructed to keep samples in personal freezers until ready to transport to lab using a provided insulated container and ice pack. Once arrived, samples were kept at −20 °C for up to a week and then moved to −80 °C until further processing. If a participant was unable to produce a sample on the requested day, they were instructed to provide the next available sample. For statistical analysis purposes, all such samples were lumped together with samples from the prior requested time point (e.g. a Tuesday sample would be lumped together with Monday samples).

Participants were instructed to continue their usual diet for week one. On week two, participants were instructed to chew every bite of food more thoroughly – approximately 30 to 40 times, or until applesauce consistency – for five days, beginning Sunday and ending on Friday when the final stool sample was collected. Diet was monitored through three dietary surveys. The Diet History Questionnaire III (DHQ3) was administered prior to the Baseline and assessed participants’ eating habits over the past month. The Automated Self-Administered 24-Hour Dietary Assessment Tool (ASA24-2018) was administered twice, once during the Baseline week and once during the Intervention Week, to provide a log of everything participants ate in the day prior to taking the test.

In some instances, a participant did not provide a stool sample on the day requested or not enough sample was present for all analyses, thus n is not always 41 for each time point. A total of 236 stool samples (average of 5.8 per person) were received. Additionally, two participants each (non-overlapping) did not complete the exit survey, DHQ3, and ASA24. In several cases, there was insufficient samples for all analyses, and one sample from Cohort 2 did not have available 16S data due to an issue during amplicon prep. Otherwise, no samples were intentionally excluded from analysis.

### Particle size measurement by laser diffraction

Stool slurries were created in an anaerobic chamber by combining stool samples and pre-reduced PBS at a 1:10 weight/volume ratio (e.g. 1 g stool with 10 mL PBS). This mixture was placed inside a custom filter bag with 2 mm pore size (in order to reduce the likelihood of clogging the laser diffraction instrument), and homogenized aerobically using a stomacher (Seward Stomacher 80 MicroBiomaster) set to medium for 60 s. Total time exposed to oxygen did not exceed 10 min. Slurries were aliquoted in 1 mL volumes and stored in cryovials at −80 °C prior to further processing. Due to the particulate nature of our samples, pipetting for this and other relevant steps was done using extra wide bore pipet tips with 2.03 mm wide orifice (Azer Scientific). Samples were randomized prior to analysis.

Fecal particle size (FPS) was measured by laser diffraction using a MasterSizer 3000 (Malvern). 500 µL stool slurry was combined with 4.5 mL 70% ethanol and incubated for at least 20 minutes in order to disinfect the sample. The following instrument settings were used: non-spherical particle type, 1.43 refractive index, 0.1 absorption, water as blank, 15 s red and 10 s blue measurement duration, 5-10 measurements per sample, 2-20% obscuration. A volume-weighted particle size distribution was produced for each sample, and summary metrics (10^th^ percentile, median, and 90^th^ percentile) were produced by the software for each sample. Each sample was added to approximately 90-100 mL diH2O in the dispersion unit, which mixed the sample through the instrument at approximately 1500-2000 rpm. The instrument was washed with a 10% bleach solution to disinfect after each use.

Pre-filtering at 2mm appeared to be reasonable given the very low abundance of particles larger than 1 mm (Fig. 2B). Given this filter, we considered any measurements reporting particles above this size to represent undesired particle aggregation, which tended to be visually apparent in the particle distributions as well. If all measurements for a sample had this problem, that sample was run again. Any individual measurements reporting particles > 2 mm were discarded as part of data pre-processing in R prior to statistical analysis.

### *In vitro* fermentation experiments

To analyze the effects of substrate particle size, wheat bran (Bob’s Red Mill) was ground by coffee grinder to produce a greater range of particle sizes, then fractionated by dry sieving. Glass beads (Next Advance) were chosen as a non-nutritive particle and were autoclaved prior to use. Both substrate types were weighed into tubes and subject to at least 15 minutes of UV sterilization in a biosafety cabinet in order to disinfect the particles as much as possible. The wheat bran experiment was carried out in 14 mL round-bottom tubes containing 2 mL pre-reduced *Bacteroides* minimal medium^64^ (BMM) containing 0.05% glucose as the sole carbon source, plus 20 mg wheat bran per tube. The glass bead experiment was carried out in 5 mL tubes containing 2 mL pre-reduced BMM with 5% glucose as the sole carbon source, plus 0-500 mg glass beads. Overnight cultures of stool-derived microbial communities previously frozen as glycerol stocks were grown in mGAM media^65^, and this culture was used to inoculate samples at 1% by volume. Cultures were incubated 24 hours at 37 °C anaerobically.

For the cross-inoculation experiment, three different Cohort 1 stool samples, selected for having higher median FPS (ZR77, PE27, and WH89 day −2 samples) were combined at 3 mL fecal slurry plus 27 mL 70% ethanol, and incubated for 90 min to disinfect. Samples were then spun down 10 min at 2000 × *g*, supernatant was discarded and samples were resuspended in 3 mL PBS. To wash residual ethanol, samples were spun down again and resuspended in fresh pre-reduced PBS. One of ten overnight cultures derived from unique stool donors from Cohort 2 was then used to inoculate each sample at 1:100 by volume (247.5 µL slurry + 2.5 µL culture). Control culture (mGAM left overnight) was used to inoculate the control wells.

### Sample fractionation

A subset of 10 stool samples were used for detailed analysis of particles by sequencing. The samples used were from the first ten participants to enroll in Cohort 1, and in all cases the first sample provided by each participant was used. Previously frozen 1 mL fecal slurry aliquots were thawed and combined with 9 mL 70% ethanol for 20 minutes to disinfect the samples; 40 mL sterile diH_2_O was then added. Three rounds of filtration were carried out in a manner adapted from methods used in aquatic systems^66^. First, gravity filtration was one by passing the sample through a 100 μm nylon mesh filter placed in a funnel and collecting the filtrate in a beaker. The filter was rinsed with another 50 mL of sterile diH_2_O to ensure any small particles had passed through. This filter was then back-rinsed with 3 mL sterile diH_2_O, and this mixture containing the retained >100 μm particles was transferred to a cryovial for storage at −80 °C.

The second and third filtration steps were done by vacuum filtration. A Buchner funnel was set up in a filter flask equipped with vacuum line, and filter paper (Whatman) was placed in the funnel and pre-wet with sterile diH_2_O. The sample was poured through with vacuum line running. As with the first filtration, filter paper was back-rinsed to remove the retained particles for the 11-100 μm and 2-11 μm fractions. The final filtrate, containing particles <2 μm in diameter, was transferred to 50 mL conical tubes and spun down for 10 min at 2,000 × *g*, and supernatant was removed prior to storage of the pellet.

### 16S rRNA gene amplicon sequencing

To assess community composition, 16S rRNA gene amplicon sequencing was performed using custom barcoded primers targeting the V4 region of the gene according to previously published protocols^67,68^. Samples newly sequenced for this study were randomized and DNA extractions were performed in tubes using the Qiagen DNeasy PowerSoil Pro DNA extraction kit (Cat no. 47016). Amplicons were cleaned (Ampure XP, Beckman Coulter, Brea, CA), quantified (QuantIT dsDNA assay kit, Invitrogen, Waltham, MA), and combined in equimolar ratios to create a sequencing pool. For any samples with post-PCR DNA concentrations insufficient to contribute enough DNA to fully balance the pool, a set volume of 7.5 μL was used. Libraries were then concentrated, gel purified, quantified again by fluorimeter, and spiked with 30% PhiX (Illumina, San Diego, CA) to improve sequencing quality. Paired-end sequencing was carried out on an Illumina MiniSeq system according to the manufacturer’s instructions using a 300-cycle Mid or High kit (Illumina, San Diego, CA, USA), depending on the number of samples in each pool.

Initial processing of raw sequence data involved custom scripts to create FASTQ files using bcl2fastq v2.20, remove primers using trimmomatic v0.36, and sync barcodes. QIIME2 was used to demultiplex sequenced samples^69^, and DADA2 was used to identify and quantify amplicon sequence variants (ASVs) in our dataset using version 138 of the Silva database^70^. We retained only samples with more than 5000 read counts to remove outlying samples that may have been subject to library preparation or sequencing artifacts^68^, and only retained taxa that appeared more than three times in at least ten percent of samples.

### Measurement of SCFAs

SCFAs were quantified by GC as previously described^71^. Briefly, randomized samples were acidified by adding 50 μL 6LJN HCl to lower the pH below 3. The mixture was vortexed and then centrifuged at 14,000 × *g* for 5 min at 4 °C. The resulting supernatant was filtered by transferring 750 μL to a 0.22 μm spin column filter and centrifuging again under the same conditions. The resulting filtrate was then transferred to a glass autosampler vial (VWR, part 66009-882). Due to the high presence of particulate matter in these samples, filters were highly prone to clogging, and multiple filtration steps were sometimes required to obtain a sufficient volume of filtrate. Filtrates were analyzed on an Agilent 7890b gas chromatograph equipped with a flame ionization detector and an Agilent HP-FFAP free fatty-acid column. A volume of 0.5LJμL of the filtrate was injected into a sampling port heated to 220 °C and equipped with a split injection liner. The column temperature was maintained at 120 °C for 1 min, ramped to 170 °C at a rate of 10 °C/min, and then maintained at 170 °C for 1 min. The helium carrier gas was run at a constant flow rate of 1 mL/min, giving an average velocity of 35 cm/s. After each sample, a 1-min post-run at 220 °C and a carrier gas flow rate of 1 mL/min was used to clear any residual sample. All C2:C5 SCFAs were identified and quantified in each sample by comparing to an eight-point standard curve ranging from 0.1 mM to 16 mM.

For statistical purposes, where a sample had no peak detected for a given SCFA, the value was set to the minimum of the lowest value detected for that compound in the run (including the standard), representing the limit of detection.

### Mouse samples

Samples from germ-free and specific-pathogen-free mice were collected from individual C57BL/6 mice (originally purchased from Jackson Laboratory and bred in-house). Germ-free mice were bred and maintained in vinyl isolators and received autoclaved food and water. Germ-free status was confirmed weekly by plating fecal samples on rich media. All mice in this study were handled in accordance with the guidelines set forth by the Duke Institutional Animal Care and Use Committee. Germ free mice have ad libitum access to autoclaved diet 5K67; specific-pathogen-free mice were given Labdiet 5053. Mice used in this study were between 13 and 29 weeks of age, and male and female mice were approximately equally represented across experiments

For the analysis of antibiotic effects, samples were from BALB/c mice (Taconic Biosciences No. BALB) from a previously published study^72^. As previously described, five cages of three mice each were used for this analysis, with diet consisting of PicoLab Rodent Diet 20 (LabDiet 5053) fed ad libitum. On days 0, 1, and 2, mice received Imipenem-Cilastatin 500mg/500mg (NDC: 63323-322-25) at 50 mg/kg body weight by oral gavage once daily. Samples were collected by cage without distinguishing individual mice.

In preparation for particle size analysis, fecal samples were combined with a volume of PBS equal to 10 times the sample weight. Since these volumes were too low for use in the stomacher as with the human samples, the mouse samples were mixed directly in 1.5 mL tubes by vortexing, and the resulting slurry was passed through a 2mm filter into a fresh tube.

### Statistics

Data analysis and visualization was done in R. When it was appropriate to compare all groups to a single intercept group (i.e. a baseline time point), linear models were performed using either the lm or lmer function in the lme4 package, with lmerTest to generate *p*-values. For linear mixed models using lmer, repeated measures sampling was handled by including participant as a random effect specified by the term (1 | participant). When multiple groups needed to be compared (e.g. clusters), an ANOVA was performed by the aov function in the stats package and, where appropriate, TukeyHSD from the same package was used as a post-hoc test to determine which specific groups differed. To calculate Spearman correlation statistics, the cor.test function in the stats package was used.

For tests of multivariate data, PERMANOVAs were performed with the adonis2 function in the vegan package, with 9999 permutations. Where appropriate, the participant term was specified for the strata argument. Differential abundance testing was done using ALDEx2^73^. The function kmeans in the stats package was used for k-means clustering of particle size distributions, after first assessing the optimal number of clusters by sum of squares and gap statistic methods (Figs. S3B-C).

## DATA AND CODE AVAILABILITY

Data and code used to generate the figures presented in this paper will be made publicly available on GitHub prior to publication. 16S rRNA gene amplicon sequence data are publicly available in the form of demultiplexed reads. Previously published data for Cohort 2 are available via the European Nucleotide Archive with accession number PRJEB47805. New data for this manuscript are available via the NCBI Sequence Read Archive (SRA) with accession numbers PRJNA902725 (Cohort 1 16S data) and PRJNA904389 (fractionation experiment 16S data).

## ACKNOWLEDGEMENTS

We would like to thank Amalia Turner, Joana Sipe, and Mark Wiesner for providing training on and access to the Malvern MasterSizer laser diffraction instrument used for particle size analysis; Otto Cordero for professional insights on the roles of particles in microbial ecosystems; Dana Hunt for guidance on particle size fractionation methods; Maria Tucker for advice on guidance for the chewing intervention; Sue Ishaq for valuable literature review recommendations; Jun Zeng for guidance on using the lyophilizer; Helene Gu for assisting with participant recruitment; Tonya Snipes and Lisa Alston-Latta for keeping our lab spaces and glassware clean; and our study volunteers for their participation. Figure 1 was created with BioRender.com.

This work was supported by National Institutes of Health grant 1R01DK116187, Office of Naval Research grant N00014-18-1-2616, Translational Research Institute through Cooperative Agreement NNX16AO69A, the Damon Runyon Cancer Research Foundation, and Burroughs Wellcome Fund Investigators in the Pathogenesis of Infectious Disease Award. This study used a high-performance computing facility partially supported by grant 2016-IDG-1013 (HARDAC+: Reproducible HPC for Next-Generation Genomics) from the North Carolina Biotechnology Center.

## REFERENCES

1. Datta, M. S., Sliwerska, E., Gore, J., Polz, M. F. & Cordero, O. X. Microbial interactions lead to rapid micro-scale successions on model marine particles. Nature communications 7, 1–7 (2016).

2. Ebrahimi, A., Goyal, A. & Cordero, O. X. Particle foraging strategies promote microbial diversity in marine environments. eLife 11, e73948 (2022).

3. Yang, X. et al. Determination of Soil Texture by Laser Diffraction Method. Soil Science Society of America Journal 79, 1556–1566 (2015).

4. Bach, E. M., Williams, R. J., Hargreaves, S. K., Yang, F. & Hofmockel, K. S. Greatest soil microbial diversity found in micro-habitats. Soil biology and Biochemistry 118, 217–226 (2018).

5. Oliver, D. M., Clegg, C. D., Heathwaite, A. L. & Haygarth, P. M. Preferential attachment of Escherichia coli to different particle size fractions of an agricultural grassland soil. Water, Air, and Soil Pollution 185, 369–375 (2007).

6. Hemkemeyer, M., Christensen, B. T., Martens, R. & Tebbe, C. C. Soil particle size fractions harbour distinct microbial communities and differ in potential for microbial mineralisation of organic pollutants. Soil Biology and Biochemistry 90, 255–265 (2015).

7. Liu, Y. et al. Differences in metabolic potential between particle-associated and free-living bacteria along Pearl River Estuary. Science of The Total Environment 728, 138856 (2020).

8. Wu, F. et al. Modulation of microbial community dynamics by spatial partitioning. Nature Chemical Biology (2022) doi:10.1038/s41589-021-00961-w.

9. Fontanez, K. M., Eppley, J. M., Samo, T. J., Karl, D. M. & DeLong, E. F. Microbial community structure and function on sinking particles in the North Pacific Subtropical Gyre. Frontiers in Microbiology 6, (2015).

10. Durkin, C. A. et al. Tracing the path of carbon export in the ocean though DNA sequencing of individual sinking particles. ISME J 16, 1896–1906 (2022).

11. Ebrahimi, A., Schwartzman, J. & Cordero, O. X. Cooperation and spatial self-organization determine rate and efficiency of particulate organic matter degradation in marine bacteria. Proceedings of the National Academy of Sciences 116, 23309–23316 (2019).

12. Matsuda, I. et al. Faecal particle size in free-ranging primates supports a ‘rumination’strategy in the proboscis monkey (Nasalis larvatus). Oecologia 174, 1127–1137 (2014).

13. Fritz, J., Hummel, J., Kienzle, E., Streich, W. J. & Clauss, M. To chew or not to chew: fecal particle size in herbivorous reptiles and mammals. Journal of Experimental Zoology Part A: Ecological Genetics and Physiology 313A, 579–586 (2010).

14. Hummel, J. et al. Differences in fecal particle size between free-ranging and captive individuals of two browser species. Zoo Biology 27, 70–77 (2008).

15. Schulz-Kornas, E., Stuhlträger, J., Clauss, M., Wittig, R. M. & Kupczik, K. Dust affects chewing efficiency and tooth wear in forest dwelling Western chimpanzees (Pan troglodytes verus). American Journal of Physical Anthropology 169, 66–77 (2019).

16. Weary, T. E., Wrangham, R. W. & Clauss, M. Applying wet sieving fecal particle size measurement to frugivores: A case study of the eastern chimpanzee (Pan troglodytes schweinfurthii). American Journal of Physical Anthropology 163, 510–518 (2017).

17. Maulfair, D. D., Fustini, M. & Heinrichs, A. J. Effect of varying total mixed ration particle size on rumen digesta and fecal particle size and digestibility in lactating dairy cows1. Journal of Dairy Science 94, 3527–3536 (2011).

18. Ishaq, S. L. et al. Pelleted-hay alfalfa feed increases sheep wether weight gain and rumen bacterial richness over loose-hay alfalfa feed. PloS one 14, e0215797 (2019).

19. Castillo-Lopez, E., Haselmann, A., Petri, R. M., Knaus, W. & Zebeli, Q. Evaluation of fecal fermentation profile and bacterial community in organically fed dairy cows consuming forage-rich diets with different particle sizes. Journal of Dairy Science 103, 8020–8033 (2020).

20. Lanyon, J. M. & Sanson, G. D. Mechanical disruption of seagrass in the digestive tract of the dugong. Journal of Zoology 270, 277–289 (2006).

21. Flint, H. J., Scott, K. P., Duncan, S. H., Louis, P. & Forano, E. Microbial degradation of complex carbohydrates in the gut. Gut Microbes 3, 289–306 (2012).

22. Lentle, R. G. & Janssen, P. W. M. Physical Aspects of the Digestion of Carbohydrate Particles. in The Physical Processes of Digestion (eds. Lentle, R. G. & Janssen, P. W. M.) 31–46 (Springer, 2011). doi:10.1007/978-1-4419-9449-3_3.

23. Mowlana, F., Heath, M. r., Van Der Bilt, A. & Van Der Glas, H. w. Assessment of chewing efficiency: a comparison of particle size distribution determined using optical scanning and sieving of almonds. Journal of Oral Rehabilitation 21, 545–551 (1994).

24. Cassady, B. A., Hollis, J. H., Fulford, A. D., Considine, R. V. & Mattes, R. D. Mastication of almonds: Effects of lipid bioaccessibility, appetite, and hormone response. American Journal of Clinical Nutrition (2009) doi:10.3945/ajcn.2008.26669.

25. Zhang, Y., Jia, J., Wang, X., Chen, J. & van der Glas, H. W. Particle size distributions following chewing: Transformation of two-dimensional outcome from optical scanning to volume outcome from sieving. Journal of Food Engineering 309, 110663 (2021).

26. Sumonsiri, P., Thongudomporn, U. & Paphangkorakit, J. Correlation between the median particle size of chewed frankfurter sausage and almonds during masticatory performance test. Journal of Oral Rehabilitation 45, 512–517 (2018).

27. Hoebler, C. et al. Physical and chemical transformations of cereal food during oral digestion in human subjects. Br J Nutr 80, 429–436 (1998).

28. Walker, A. W. et al. The species composition of the human intestinal microbiota differs between particle-associated and liquid phase communities. Environmental Microbiology 10, 3275–3283 (2008).

29. Jenkins, D. J. A. et al. The effect of wheat bran particle size on laxation and colonic fermentation. Journal of the American College of Nutrition 18, 339–345 (1999).

30. Tuncil, Y. E., Thakkar, R. D., Marcia, A. D. R., Hamaker, B. R. & Lindemann, S. R. Divergent short-chain fatty acid production and succession of colonic microbiota arise in fermentation of variously-sized wheat bran fractions. Scientific reports 8, 1–13 (2018).

31. Thakkar, R. D., Tuncil, Y. E., Hamaker, B. R. & Lindemann, S. R. Maize Bran Particle Size Governs the Community Composition and Metabolic Output of Human Gut Microbiota in in vitro Fermentations. Frontiers in Microbiology 11, 1009 (2020).

32. De Paepe, K. et al. Modification of wheat bran particle size and tissue composition affects colonisation and metabolism by human faecal microbiota. Food \& function 10, 379–396 (2019).

33. De Paepe, K., Verspreet, J., Courtin, C. M. & de Wiele, T. Microbial succession during wheat bran fermentation and colonisation by human faecal microbiota as a result of niche diversification. The ISME journal 14, 584–596 (2020).

34. Letourneau, J. et al. Ecological memory of prior nutrient exposure in the human gut microbiome. ISME J 1–12 (2022) doi:10.1038/s41396-022-01292-x.

35. Holmes, Z. C. et al. Microbiota responses to different prebiotics are conserved within individuals and associated with habitual fiber intake. Microbiome 10, 114 (2022).

36. Rose, C., Parker, A., Jefferson, B. & Cartmell, E. The characterization of feces and urine: A review of the literature to inform advanced treatment technology. Critical Reviews in Environmental Science and Technology (2015) doi:10.1080/10643389.2014.1000761.

37. Petrone, B. L. et al. Diversity of plant DNA in stool is linked to dietary quality, age, and household income. Proceedings of the National Academy of Sciences 120, e2304441120 (2023).

38. Miquel-Kergoat, S., Azais-Braesco, V., Burton-Freeman, B. & Hetherington, M. M. Effects of chewing on appetite, food intake and gut hormones: A systematic review and meta-analysis. Physiology \& behavior 151, 88–96 (2015).

39. Naumova, E. I. et al. Particle size reduction along the digestive tract of fat sand rats (Psammomys obesus) fed four chenopods. J Comp Physiol B 191, 831–841 (2021).

40. Lewis, S. J. & Heaton, K. W. Stool Form Scale as a Useful Guide to Intestinal Transit Time. Scandinavian Journal of Gastroenterology 32, 920–924 (1997).

41. Procházková, N. et al. Advancing human gut microbiota research by considering gut transit time. Gut gutjnl-2022-328166 (2022) doi:10.1136/gutjnl-2022-328166.

42. Ruppin, H., Bar-Meir, S., Soergel, K. H., Wood, C. M. & Schmitt, M. G. Absorption of Short-Chain Fatty Acids by the Colon. Gastroenterology 78, 1500–1507 (1980).

43. Gophna, U., Konikoff, T. & Nielsen, H. B. *Oscillospira* and related bacteria - From metagenomic species to metabolic features: Metabolism of Oscillospira. Environ Microbiol 19, 835–841 (2017).

44. Romano, S. et al. Meta-analysis of the Parkinson’s disease gut microbiome suggests alterations linked to intestinal inflammation. npj Parkinsons Dis. 7, 1–13 (2021).

45. McCulloch, J. A. & Trinchieri, G. Gut bacteria enable prostate cancer growth. Science 374, 154–155 (2021).

46. Wichmann, A. et al. Microbial Modulation of Energy Availability in the Colon Regulates Intestinal Transit. Cell Host & Microbe 14, 582–590 (2013).

47. Boekhorst, J. et al. Stool energy density is positively correlated to intestinal transit time and related to microbial enterotypes. Microbiome 10, 223 (2022).

48. Carmody, R. N., Weintraub, G. S. & Wrangham, R. W. Energetic consequences of thermal and nonthermal food processing. Proceedings of the National Academy of Sciences 108, 19199–19203 (2011).

49. Lucas, P. W. et al. Measuring the Toughness of Primate Foods and its Ecological Value. Int J Primatol 33, 598–610 (2012).

50. Zink, K. D., Lieberman, D. E. & Lucas, P. W. Food material properties and early hominin processing techniques. Journal of Human Evolution 77, 155–166 (2014).

51. Hardy, K., Brand-Miller, J., Brown, K. D., Thomas, M. G. & Copeland, L. The Importance of Dietary Carbohydrate in Human Evolution. The Quarterly Review of Biology 90, 251–268 (2015).

52. Wright, B. W. et al. It’s Tough Out There: Variation in the Toughness of Ingested Leaves and Feeding Behavior Among Four Colobinae in Vietnam. Int J Primatol 29, 1455–1466 (2008).

53. Hebelstrup, K. H. Differences in nutritional quality between wild and domesticated forms of barley and emmer wheat. Plant Science 256, 1–4 (2017).

54. Cordain, L. et al. Origins and evolution of the Western diet: health implications for the 21st century1,2. The American Journal of Clinical Nutrition 81, 341–354 (2005).

55. Fritz, J. et al. Comparative chewing efficiency in mammalian herbivores. Oikos 118, 1623–1632 (2009).

56. Patricia, J. J. & Dhamoon, A. S. Physiology, Digestion. (2019).

57. Reese, A. T. et al. Antibiotic-induced changes in the microbiota disrupt redox dynamics in the gut. eLife 7, e35987 (2018).

58. Kumari, M. & Kozyrskyj, A. L. Gut microbial metabolism defines host metabolism: an emerging perspective in obesity and allergic inflammation. Obesity Reviews 18, 18–31 (2017).

59. Moco, S. et al. Systems Biology Approaches for Inflammatory Bowel Disease: Emphasis on Gut Microbial Metabolism. Inflammatory Bowel Diseases 20, 2104–2114 (2014).

60. Wissel, E. F. & Smith, L. K. Inter-individual variation shapes the human microbiome. Behav Brain Sci 42, (2019).

61. De Paepe, K., Kerckhof, F.-M., Verspreet, J., Courtin, C. M. & Van de Wiele, T. Inter-individual differences determine the outcome of wheat bran colonization by the human gut microbiome. Environmental Microbiology 19, 3251–3267 (2017).

62. Welker, T. L., Overturf, K. & Barrows, F. Development and Evaluation of a Volumetric Quantification Method for Fecal Particle Size Classification in Rainbow Trout Fed Different Diets. North Am J Aquaculture 82, 159–168 (2020).

63. Turnbaugh, P. J. et al. An obesity-associated gut microbiome with increased capacity for energy harvest. Nature 444, 1027–1031 (2006).

64. Villa, M. M. et al. Interindividual Variation in Dietary Carbohydrate Metabolism by Gut Bacteria Revealed with Droplet Microfluidic Culture. mSystems 5, e00864–19 (2020).

65. Rettedal, E. A., Gumpert, H. & Sommer, M. O. A. Cultivation-based multiplex phenotyping of human gut microbiota allows targeted recovery of previously uncultured bacteria. Nat Commun 5, 1–9 (2014).

66. Hunt, D. E. et al. Resource partitioning and sympatric differentiation among closely related bacterioplankton. Science 320, 1081–1085 (2008).

67. Caporaso, J. G. et al. Ultra-high-throughput microbial community analysis on the Illumina HiSeq and MiSeq platforms. ISME J 6, 1621–1624 (2012).

68. Silverman, J. D., Durand, H. K., Bloom, R. J., Mukherjee, S. & David, L. A. Dynamic linear models guide design and analysis of microbiota studies within artificial human guts. Microbiome 6, 1–20 (2018).

69. Caporaso, J. G. et al. QIIME allows analysis of high-throughput community sequencing data. Nature Methods (2010) doi:10.1038/nmeth.f.303.

70. Callahan, B. J. et al. DADA2: High-resolution sample inference from Illumina amplicon data. Nat Methods 13, 581–583 (2016).

71. Holmes, Z. C. et al. Short-chain fatty acid production by gut microbiota from children with obesity differs according to prebiotic choice and bacterial community composition. mBio 11, e00914–20 (2020).

72. Holmes, Z. C. et al. Prebiotic galactooligosaccharides interact with mouse gut microbiota to attenuate acute graft-versus-host disease. Blood (2022) 10.1182/blood.2021015178.

73. Fernandes, A. D., Macklaim, J. M., Linn, T. G., Reid, G. & Gloor, G. B. ANOVA-Like Differential Expression (ALDEx) Analysis for Mixed Population RNA-Seq. PLoS One 8, e67019 (2013).

