## Supplementary Figures S1-S13 for "Interplay between particle size and microbial ecology in the gut microbiome"

**This file includes:**

Figures S1 to S13

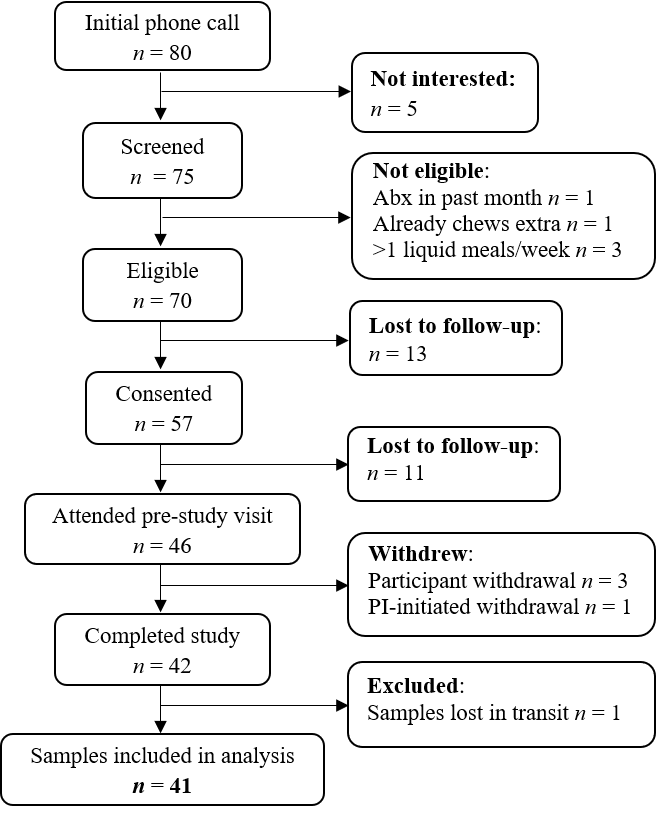

**Figure S1. Recruitment and enrollment statistics for Cohort 1.**

Analytic sample size flowchart in the style of Strengthening The Organization and Reporting of Microbiome Studies (STORMS).

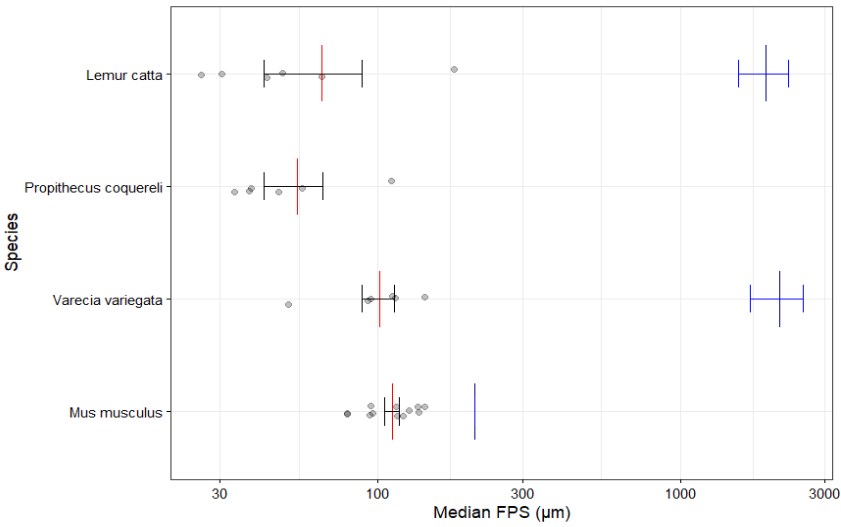

**Figure S2. Comparison of measured FPS to previously reported values across species.**

Median FPS shown on log scale for three species of lemur: *Lemur catta* (*n* = 6), *Propithecus coquereli*, (*n* = 6), and *Varecia variegata* (*n* = 6), as well as specific pathogen-free (SPF) mice (*n* = 10). Previously reported values from Fritz et al. (*J. Anim. Physiol. Anim. Nutr.*, 2009) mean and standard error shown (*Lemur catta* *n* = 3, *Propithecus coquereli* no data, *Varecia variegata n* = 4, and *Mus musculus* *n* = 1).

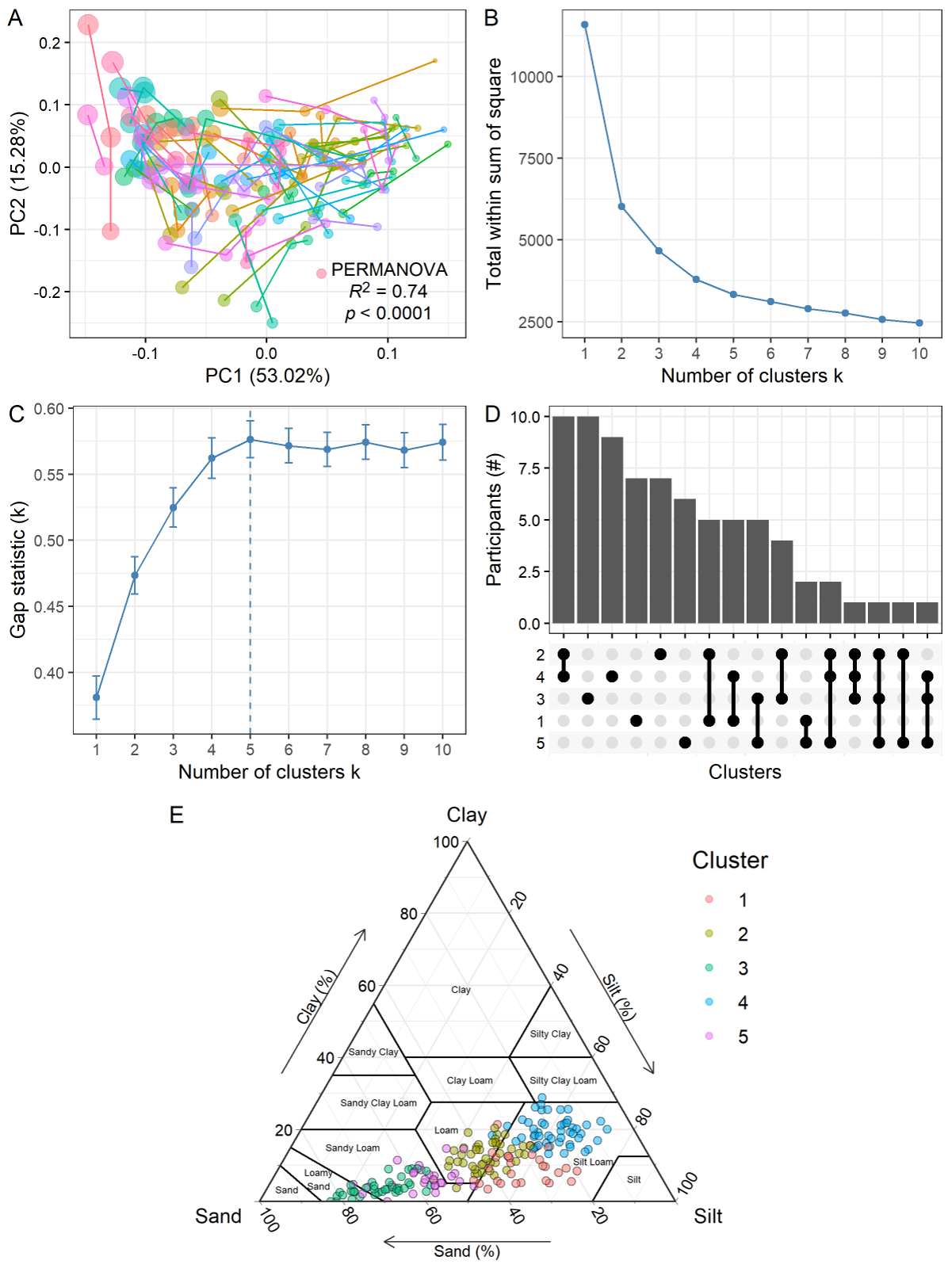

**Figure S3. Analysis of baseline particle size distributions.**

**A**, Principal component analysis of baseline sample particle size distributions, with points colored and connected by participant. Point size corresponds to median FPS for that sample. PERMANOVA statistic calculated on participant. (*n* = 186 samples from 76 participants; 41 in Cohort 1 and 35 in Cohort 2.) **B-C**, Total within sum of square (**B**) and gap statistic (**C**) plots used to determine the optimal number of clusters for k-means clustering. **D**, Upset plot showing the frequency of samples for each participant appearing in each cluster or combination of clusters (i.e. 10 participants had samples in both clusters 2 and 4, 10 participants had samples only in cluster 3, etc.). **E**, Ternary plot showing baseline samples on a USDA soil textural classification diagram, where sand is the largest particle size (50-2000 µm), silt is intermediate (2-50 µm), and clay is the smallest (< 2 µm).

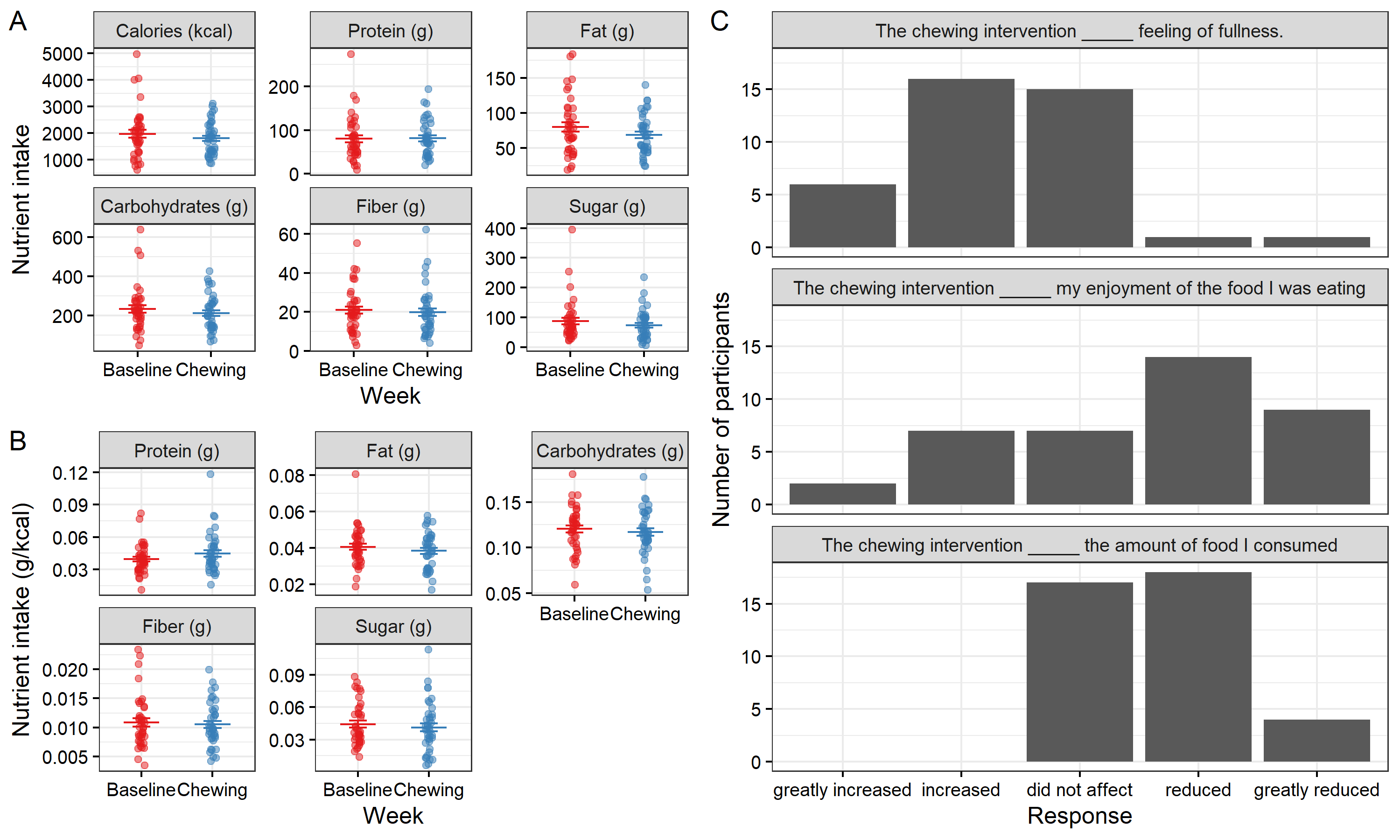

**Figure S4. Chewing study diet and exit survey data.**

**A-B**, Macronutrient intake reported on ASA24 by week in terms of raw values (**A**) or proportions relative to total kcal (**B**). Mean and standard error plotted. Linear mixed model (week as fixed effect, participant as random effect) *p* > 0.05 in all cases. **C**, Frequencies of responses to exit survey questions about chewing intervention. **A-C**, (*n* = 39 participants).

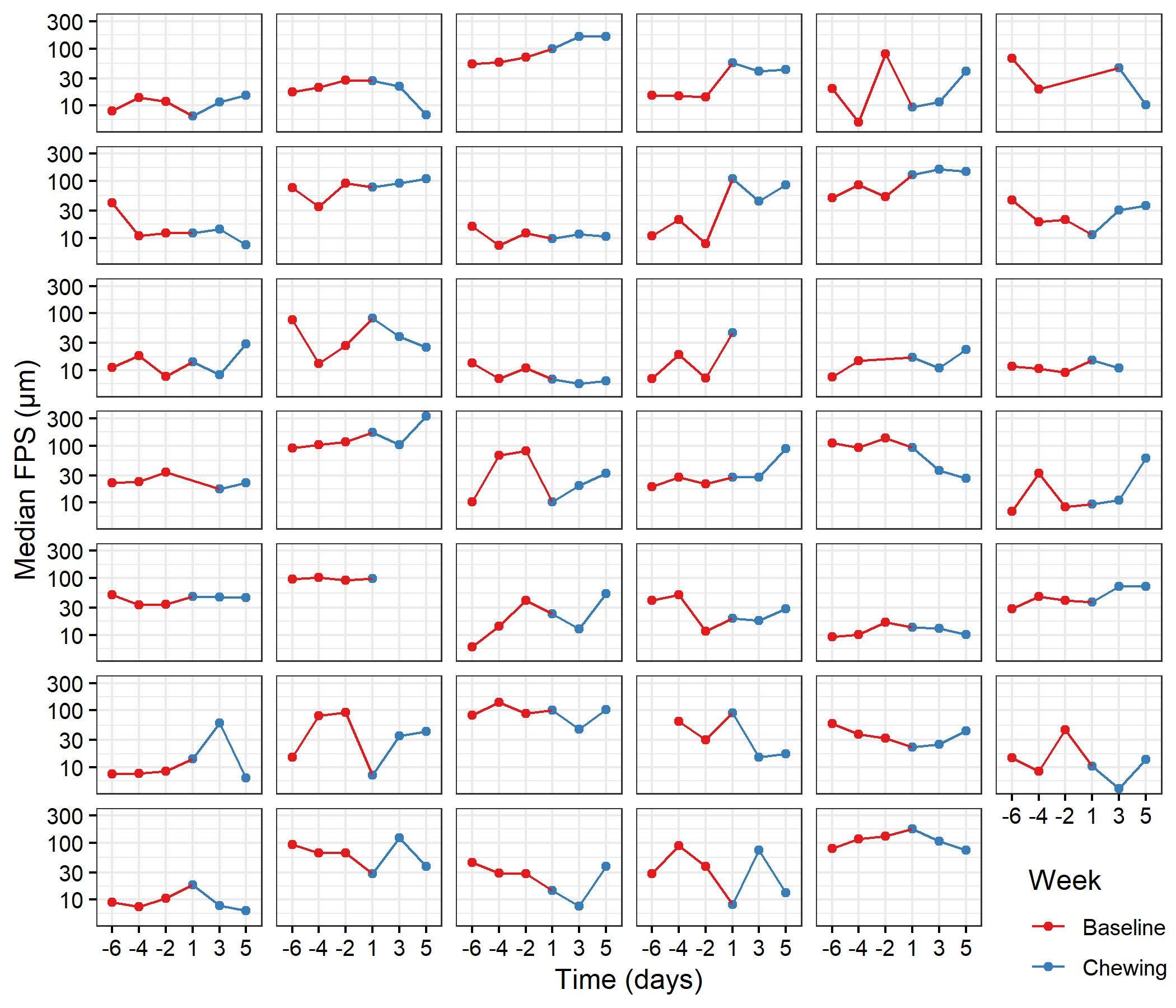

**Figure S5. Trends in median FPS by individual.**

Plots of median FPS over time for each individual in Cohort 1, where red points are from the baseline week and blue points are from the chewing week.

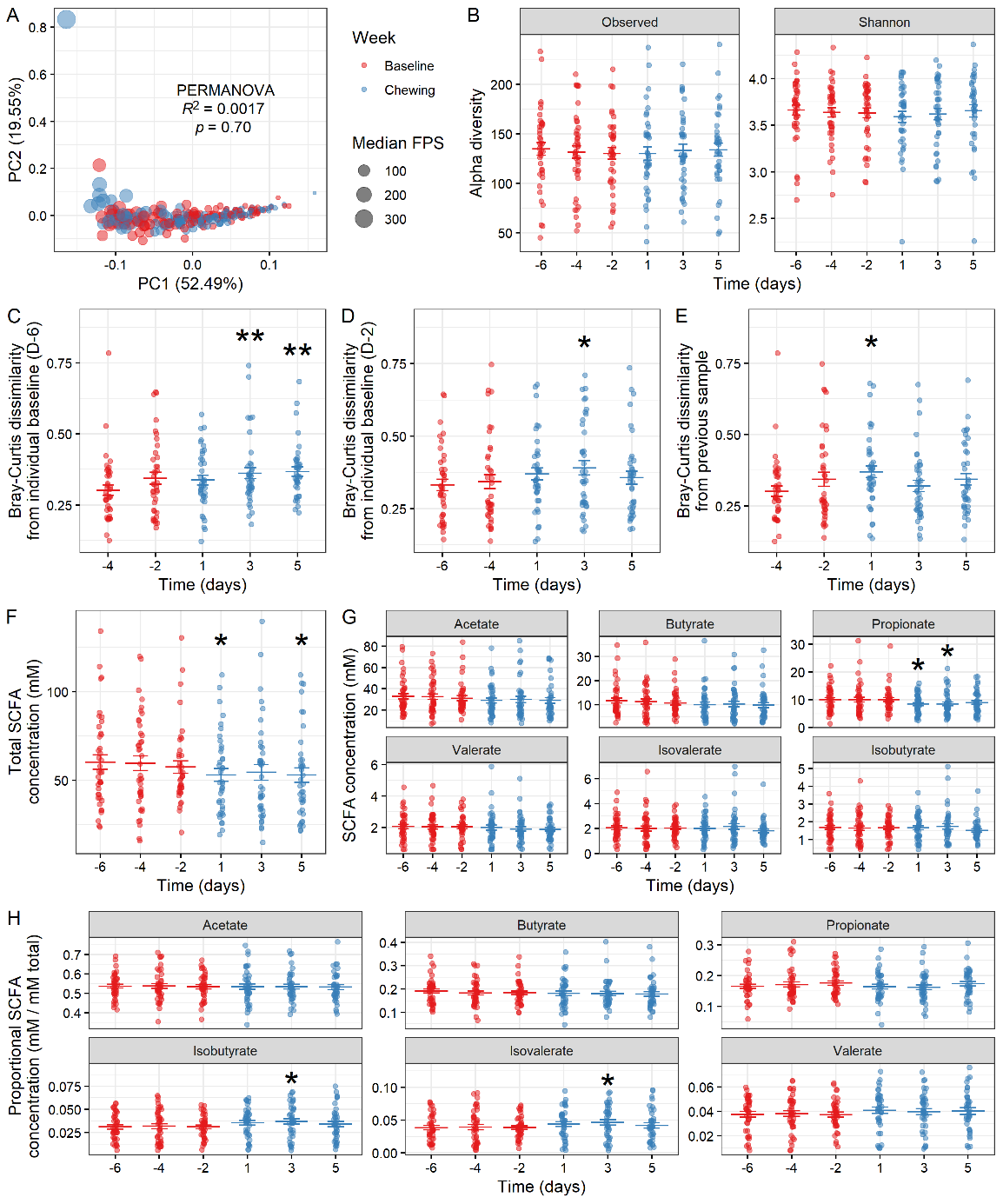

**Figure S6. Chewing study microbiome composition and SCFA data.**

**A**, PCA plot of particle size distributions by week, with PERMANOVA on week (participant as strata) shown. (*n* = 236 samples from 41 participants). **B**, Alpha diversity by observed ASVs and Shannon index by time point. Linear mixed model (day as fixed effect, participant as random effect), with day -6 as intercept, *p* > 0.05; for week as fixed effect, *p* = 0.53 for observed ASVs and *p* = 0.21 for Shannon. **C-E**, Bray-Curtis dissimilarity relative to each participant’s first sample provided (**C**), the day -2 sample (**D**), or the previous sample (**E**). Linear mixed model (day as fixed effect, participant as random effect), with earliest time point as intercept, shown; for week as fixed effect, *p* = 0.026 (**C**), *p* = 0.0133 (**D**), and *p* = 0.17 (**E**). **F-H**, Total (**F**) and individual (**G**) SCFA concentrations or proportions (**H**) by time point. Linear mixed models (day as categorical fixed effect, participant as random effect), with day -6 as intercept, shown; for week as fixed effect, *p* < 0.05 for acetate, propionate, and valerate concentrations (**F**), and *p* < 0.05 for isobutyrate and isovalerate proportions (**H**). **B-H**, Mean and standard error plotted. (*n* = 41 participants). * *p* < 0.05, ** *p* < 0.01, *** *p* < 0.001.

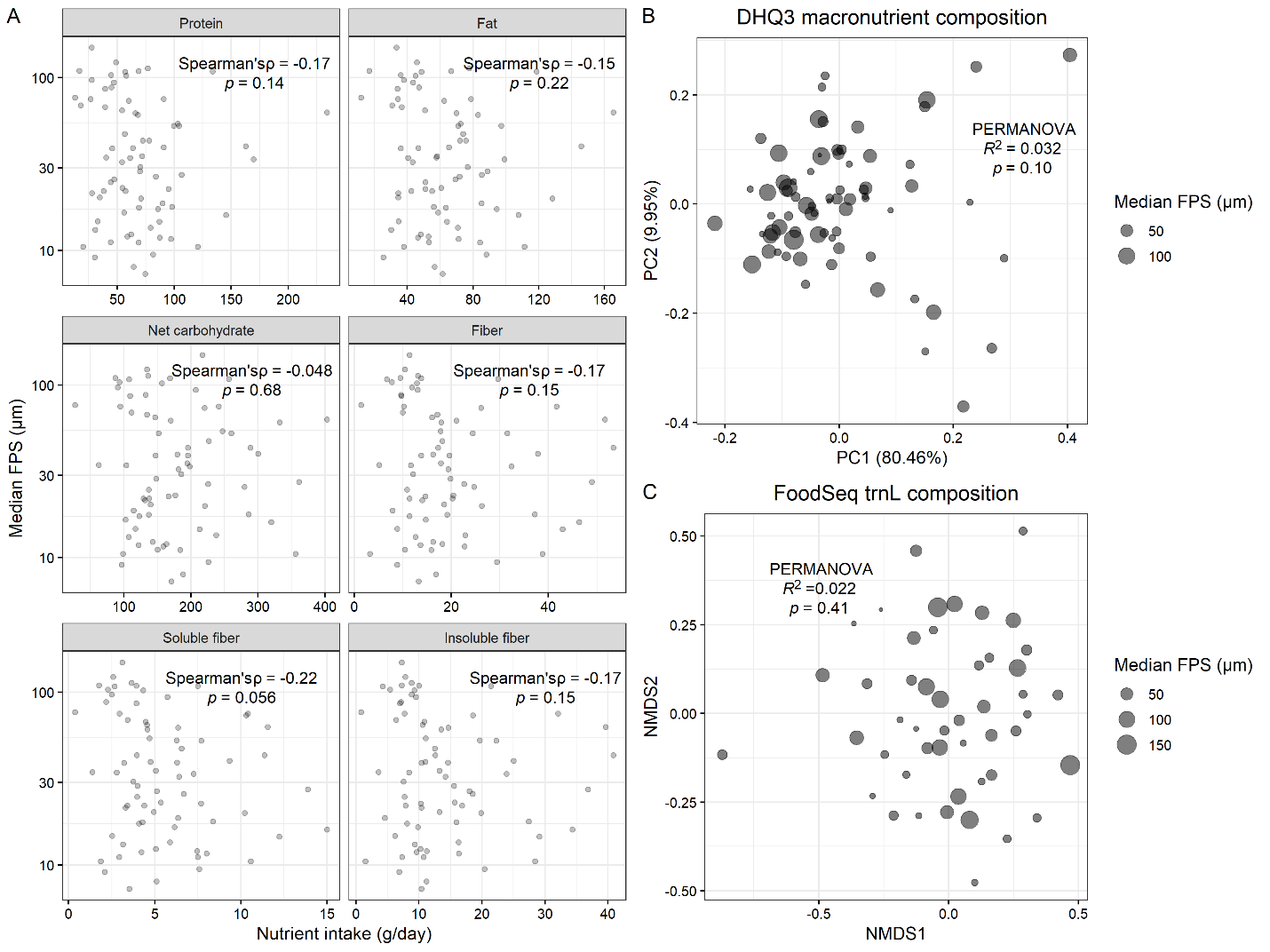

**Figure S7. Relationship of FPS with diet.**

**A**, Relationship between habitual intake of macronutrients as reported by DHQ3, with a focus on dietary fiber, and median FPS. Results of Spearman correlation test shown. **B**, Data from (**A**) ordinated onto PCA space, with results of PERMANOVA by median FPS shown. **A-B**, (*n* = 73 participants.) **C,** NMDS plot of food DNA by trnL sequencing using FoodSeq, with PERMANOVA by median FPS with participant as strata shown (Cohort 2 only; *n* = 35 participants).

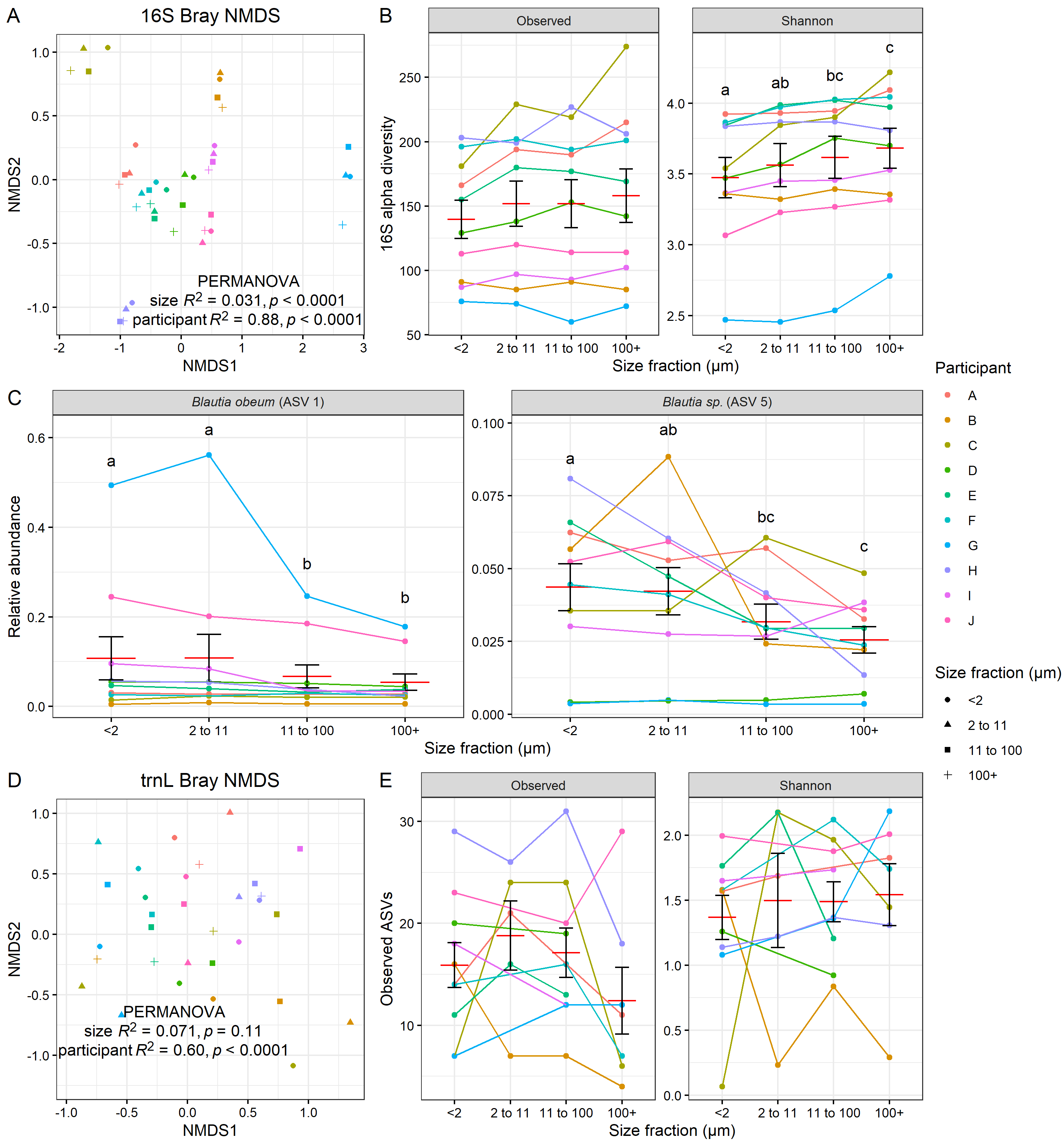

**Figure S8. Sequencing of particle fractions of ten samples.**

**A**, NMDS ordination plot of 16S rRNA gene amplicon sequencing of ten fractionated stool samples from different participants. Results of PERMANOVA (size + participant) shown. **B**, 16S alpha diversity by particle size fraction. ANOVA (size + participant) for Observed: size *p* = 0.050, participant *p* = 8.7 × 10^-16^; for Shannon: size *p* = 0.00025, donor *p* < 2 × 10^-16^; letters indicate significantly different (*p* < 0.05) groups by Tukey HSD test. **C**, 16S ASVs identified as significantly different by size fraction by ALDEx2 GLM (size + participant); letters indicate significantly different (*p* < 0.05) groups by Tukey HSD test. **D**, NMDS ordination plot of trnL diet metabarcoding sequencing of. Results of PERMANOVA (size + participant) shown. **E**, trnL alpha diversity by particle size fraction. ANOVA (size + participant) for Observed: size *p* = 0.054, participant *p* = 0.22; for Shannon: size *p* = 0.13, donor *p* = 0.95. **A-E**, (*n* = 10 participants.)

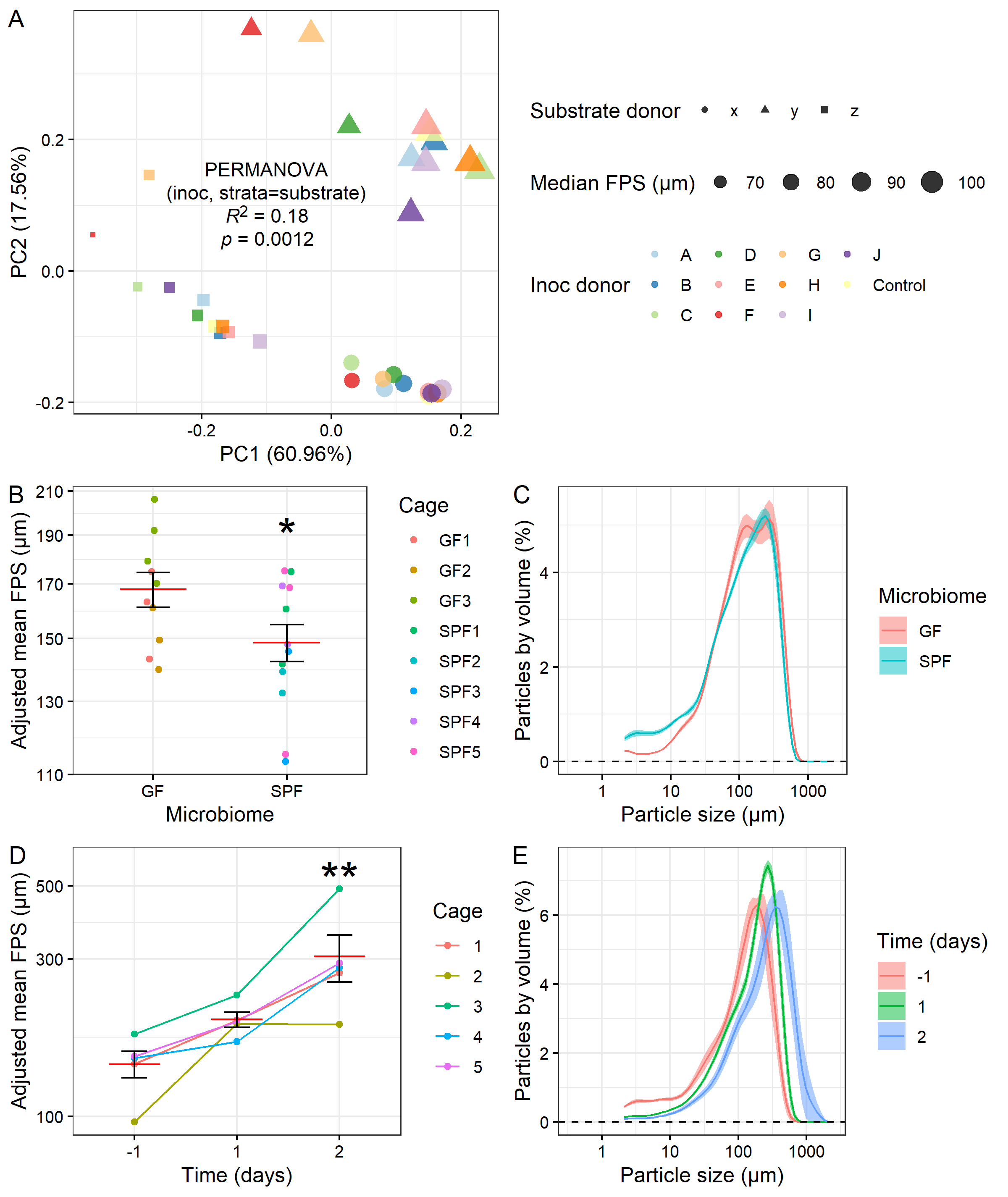

**Figure S9. Role of microbiome in fecal particle breakdown.**

**A**, PCA plot of post-incubation particle size distributions for *in vitro* cross-inoculation experiment. Stool samples from three donors were disinfected by incubation in ethanol, then inoculated with cultures from one of ten additional stool donors, or no-inoculation control. Result of PERMANOVA on inoculation donor with substrate donor as strata shown. **B-D**, Data for mouse experiments shown in Fig. 5, recomputed to omit particles < 2 µm. **B**, Mean FPS in germ-free (GF) and specific-pathogen-free (SPF) mice collected from individual mice. (Linear model with GF as intercept; *n* = 10 GF mice and 12 SPF mice.) **C**, Particle size distributions for (**B**). **D**, Mean FPS for mouse fecal samples collected by cage, with imipenem antibiotic treatment begun on day 0. Linear mixed model (day as categorical fixed effect, cage as random effect) with D-1 as intercept shown. (*n* = 5 cages.) **E**, Particle size distributions for (**D**). **B-D**, Mean and standard error plotted. * *p* < 0.05, ** *p* < 0.01, *** *p* < 0.001.

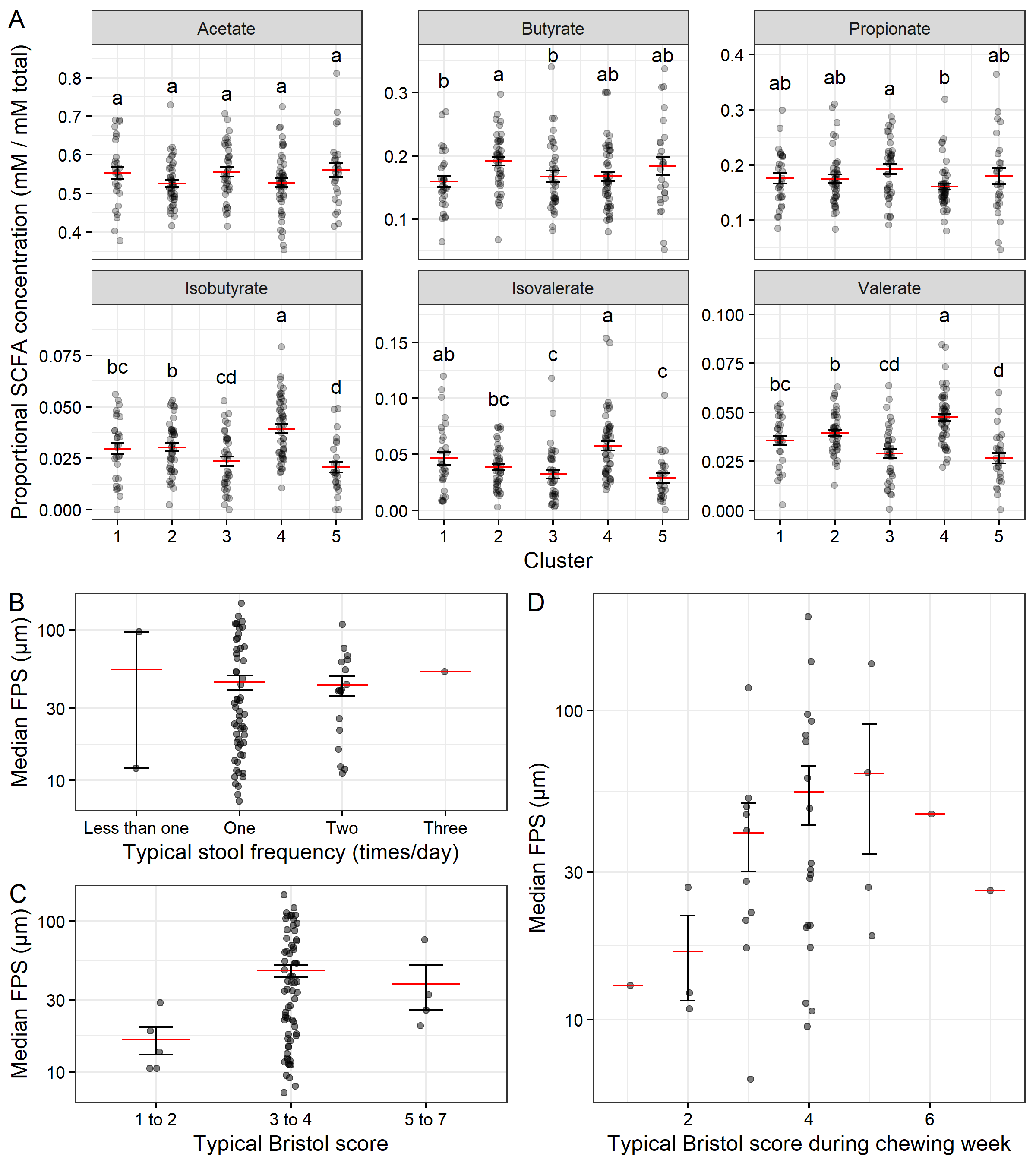

**Figure S10. Relationships of FPS with SCFA proportions and stool statistics.**

**A,** Proportional SCFA concentrations for all baseline samples by cluster. ANOVAs (cluster + participant) significant for both variables for total all individual SCFAs; letters indicate significantly different groups by Tukey HSD test. (*n* = 185 samples.) **B-C,** Self-reported typical stooling frequencies **(B)** and Bristol stool scores **(C**) plotted against average median FPS for each participant. ANOVA *p* > 0.05 for both variables (*p* = 0.15 for Bristol score and *p* = 0.97 for frequency; *n* = 76 participants.) **D,** Exit survey responses to the question “What type of stool did you most often experience over the past week?” in Cohort 1, plotted against the average median FPS within that week for each participant. (*n* = 39 participants.) **A-D,** Mean and standard error plotted.

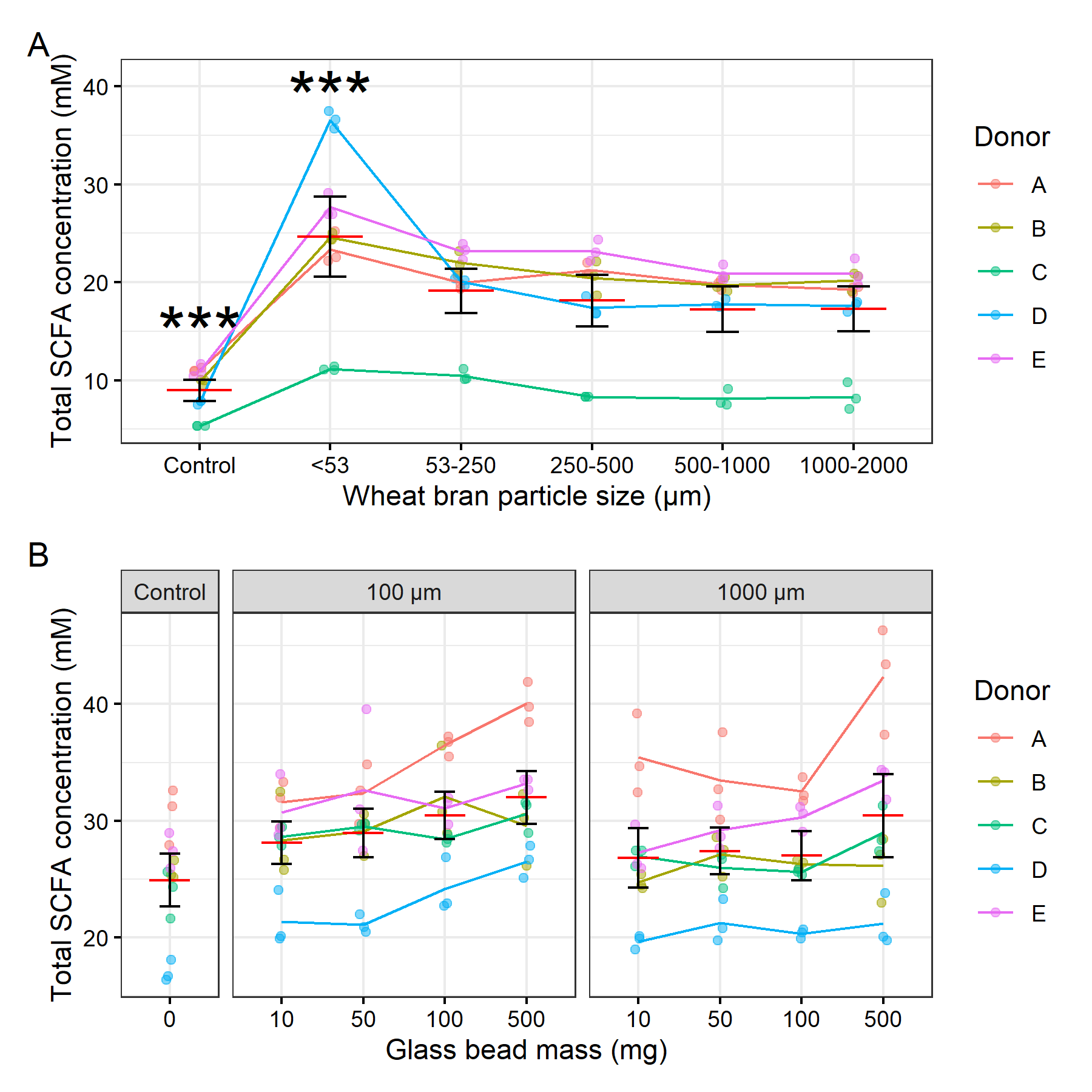

**Figure S11. Substrate particle size influences gut microbial metabolism in vitro.**

**A,** Total SCFA concentration after 24-hour incubation of stool-derived microbial communities with wheat bran particles of different sizes. Linear mixed model (size as fixed effect, donor as random effect) with 1000-2000 as intercept shown. (*n* = 5 donors with 3 technical replicates each.) **B,** Total SCFA concentration after 24-hour incubation of stool-derived microbial communities with small or large glass beads added at different masses. Linear mixed model (size:mass as fixed effects, donor as random effect), excluding control, with 100 µm glass beads and 10 mg bead mass as intercept conditions, gives size *p* = 0.0014, mass *p* = 0.00013, size:mass p = 0.83. (*n* = 5 donors with 3 technical replicates each.) Mean and standard error plotted. * *p* < 0.05, ** *p* < 0.01, *** *p* < 0.001.

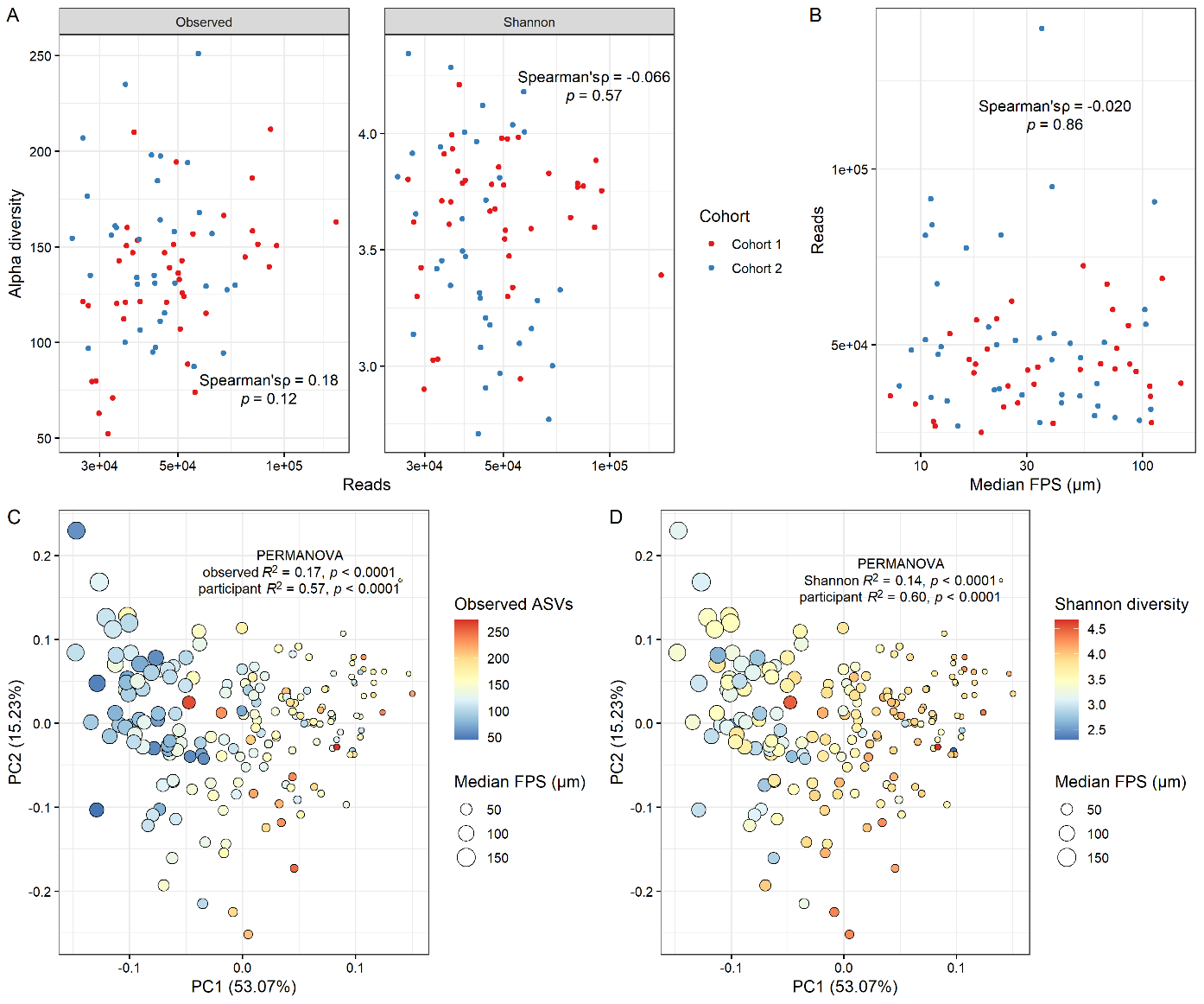

**Figure S12. FPS and alpha diversity.**

**A**, Scatterplot of read depth averaged within each participant and alpha diversity by observed ASVs and Shannon index. Spearman correlation test statistics shown. (*n* = 76 participants; 41 in Cohort 1 and 35 in Cohort 2.) **B**, Scatterplot of median FPS averaged within each participant and read depth. Spearman correlation test statistics shown. **C-D**, PCA of particle size distributions colored by alpha diversity using observed ASVs (**C**) or Shannon index (**D**), with results of PERMANOVA (diversity + participant) shown. (*n* = 185 samples.)

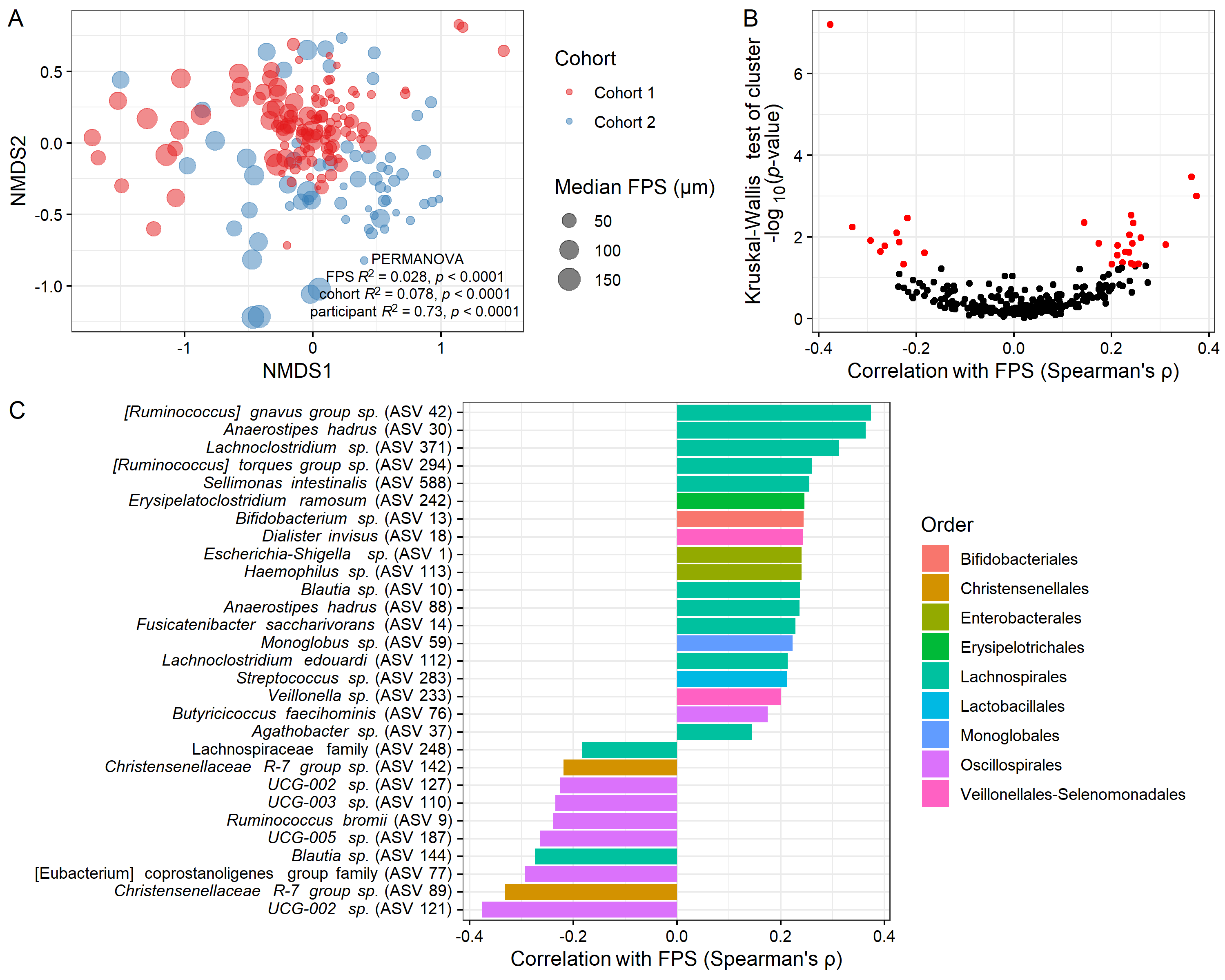

**Figure S13. FPS and community composition.**

**A**, NMDS ordination plot equivalent to Fig. 2C, colored by cohort, with results of PERMANOVA (cluster + cohort + participant) shown. (*n* = 185 samples.) **B**, Volcano plot depicting results of ALDEx2 Kruskal-Wallis test on particle size distribution cluster, with ASVs found to be significantly (FDR-corrected *p* < 0.05) differentially abundant by cluster shown in red. Each point is an ASV (298 tested, 29 significant), and each ASV’s Spearman correlation with FPS is shown on the x-axis as an effect size metric. **C**, Identities and Spearman correlations of ASVs found to be significantly differentially abundant by cluster (red points in (**B**)), colored by order.
